# Miro GTPase domains regulate assembly of the mitochondrial motor-adaptor complex

**DOI:** 10.1101/2022.05.13.491728

**Authors:** Kayla Davis, Himanish Basu, Ethan Shurberg, Thomas L. Schwarz

**Affiliations:** Kirby Neurobiology Center, Boston Children’s Hospital, Boston, MA 02115, USA; Division of Medical Sciences, Harvard Medical School, Boston, MA 02115, USA; Department of Immunology, Harvard Medical School, Boston, MA, 02115; Department of Neurobiology, Harvard Medical School, Boston, MA 02115, USA

**Keywords:** Mitochondria, motor-adaptor complex, Miro1, Miro2, RHOT1, RHOT2, TRAK1, P135, KIF5, GTPase, trafficking

## Abstract

Mitochondrial transport relies on a motor-adaptor complex containing Miro1, a mitochondrial outer membrane protein with two GTPase domains, as well as TRAK1/2, kinesin-1, and dynein. Using a peroxisome-directed Miro1, we quantified the ability of GTPase mutations to influence peroxisomal recruitment of complex components. Miro1 whose N-GTPase is locked in the GDP-state doesn’t recruit TRAK1/2, kinesin or P135 to peroxisomes whereas the GTP-state does. Miro1 C-GTPase mutations have little influence on complex recruitment. Though Miro2 is thought to support mitochondrial motility, peroxisome-directed Miro2 did not recruit the other complex components regardless of the state of its GTPase domains. Neurons expressing peroxisomal Miro1 with the GTP-state form of the N-GTPase had markedly increased peroxisomal transport to growth cones while the GDP-state caused their retention in the soma. Thus, the N-GTPase of Miro1 is critical for regulating Miro1’s interaction with the other components of the motor-adaptor complex and thereby for regulating mitochondrial motility.

**Summary:** A Miro-containing complex mediates mitochondrial motility. Relocalizing Miro1 and 2 to peroxisomes and systematically manipulating each GTPase domain of Miro revealed the importance of the N-terminal GTPase domain of Miro1 for governing interaction with TRAK proteins, motors, and transport.

## Introduction

Mitochondria move in order to distribute themselves and ensure that they can efficiently respond to metabolic demands and accomplish local calcium buffering and ROS signaling (Emptage et al., 2001; MacAskill et al., 2009a; Marchi et al., 2012; Mattson et al., 2008; Mochida et al., 2008). The proper distribution of mitochondria is critical for their inheritance during cell division and their contacts with other organelles(Chung et al., 2016; Kanfer et al., 2015; Modi et al., 2019). In animal cells, mitochondria are trafficked along microtubules by a motor-adaptor protein complex (Brickley et al., 2005; Glater et al., 2006; Hirokawa et al., 1991; Pilling et al., 2006; Stowers et al., 2002; van Spronsen et al., 2013). Disrupting this trafficking can lead to neurodegeneration (Hardy, 2010; Liu et al., 2012; Morotz et al., 2012; Nguyen et al., 2014; Wang et al., 2011; Zhang et al., 2015). The motor-adaptor complex includes Miro, an outer mitochondrial membrane protein and Rho-like GTPase, TRAK, a motor-adaptor protein, and two microtubule-based motors (Brickley and Stephenson, 2011; Fransson et al., 2003; Fransson et al., 2006; Glater et al., 2006; Stowers et al., 2002). The molecular motor kinesin-1 (KIF5) transports mitochondria towards the plus-end of microtubules, and the dynein-dynactin complex transports mitochondria towards the minus-end of microtubules (Chada and Hollenbeck, 2003; Drerup et al., 2017; Glater et al., 2006; Hurd and Saxton, 1996; Pilling et al., 2006; Tanaka et al., 1998; Vale et al., 1985). Additional components of the motor-adaptor complex have been found. Syntaphilin (SNPH) anchors mitochondria to microtubules, FHL2 anchors mitochondria to actin, and Myo19 drives short-range movements on actin filaments (Basu et al., 2021; Bocanegra et al., 2020; Chen and Sheng, 2013; Kang et al., 2008; Lopez-Domenech et al., 2018; Lopez-Domenech et al., 2021; Norkett et al., 2020; Oeding et al., 2018; Quintero et al., 2009; Seo et al., 2018). DISC1, O-GlcNAc transferase, and metaxins have also been implicated in the function and regulation of the motor-adaptor complex (Ogawa et al., 2014; Pekkurnaz et al., 2014; Zhao et al., 2021).

Mice and humans have two Miro genes, Miro1 and Miro2 (also called RhoT1 and 2), but their functional differences are not well understood (Fransson et al., 2003). Miro1 plays a central role in mitochondrial dynamics and is critical for mammalian development. Miro1 knockout is lethal in mice and decreases mitochondrial trafficking in somatic cells and neurons (Babic et al., 2015; Lopez-Domenech et al., 2018; Lopez-Domenech et al., 2016; Nguyen et al., 2014). Miro1 also participates in mitochondrial turnover, mitochondria-ER contact sites, and mitochondrial fission and fusion (Bocanegra et al., 2020; Lee et al., 2018; Lopez-Domenech et al., 2018; Lopez-Domenech et al., 2021; Misko et al., 2010; Modi et al., 2019; Shlevkov et al., 2016; Wang et al., 2011). There are four splicing variants of Miro1; some are known to localize to peroxisomes in at least some cell types or upon over-expression (Castro and Schrader, 2018; Covill-Cooke et al., 2020; Okumoto et al., 2018).

Miro has a C-terminal transmembrane domain that anchors the protein to mitochondria and two GTPase domains, one at the N-terminal and one near the C-terminal; these two GTPase domains are the focus of the current manuscript. These GTPase domains are separated by two pairs of EF-hands (Fransson et al., 2003; Klosowiak et al., 2013; Smith et al., 2020) which cause mitochondrial movement to stop when cytosolic Ca^2+^ increases (Macaskill et al., 2009b; Saotome et al., 2008; Wang et al., 2011). The functional significance of the GTPase domains has been harder to establish. Overexpressing Miro1 N-GTPase locked in a constitutively active, GTP-bound state (P13V) alters mitochondrial distribution, while overexpressing mutations that place the N-GTPase in a constitutively inactive, GDP-bound state (T18N) decreases mitochondrial trafficking and alters their distribution (Babic et al., 2015; Fransson et al., 2003; Fransson et al., 2006). A Vibrio cholerae-derived protein that activates the GTPase activity of Miro converts Miro to the GDP-bound state and thereby alters mitochondrial distribution (Suzuki et al., 2014). Mitochondria with the T18N Miro1 mutation prevent the binding of Myo19, CENP-F and DISC1 (Kanfer et al., 2015; Norkett et al., 2020; Oeding et al., 2018). Although these studies of the GTPase domains imply a regulatory role, the mechanism by which they alter mitochondrial distribution is less clear, particularly as concerns their transport by microtubule-based motors.

Determining the how the GTPase domains influence transport by overexpression of mutant forms has been confounded by three issues: 1) the presence of the endogenous Miro1 and Miro2 on mitochondria, 2) the deleterious effects of grossly disturbing mitochondrial distribution and the other functions of Miro, and 3) the presence of other TRAK-binding proteins on mitochondria that may provide a parallel or alternative means of supporting transport (Lopez-Domenech et al., 2016; Misko et al., 2010). The formation of dimers between endogenous wildtype Miro and an expressed mutant, for example, can mask the consequences of the mutation on the assembly of the motor complex. The endogenous Miro or Miro-independent paths may also continue to mediate transport even when a completely inactive Miro is also expressed. To study the function of the GTPase domains with an independent approach and without these confounding factors, we’ve misdirected Miro transgenes to peroxisomes by using constructs that lack the mitochondria-targeting transmembrane domain and instead have a PEX3 peroxisomal-targeting sequence. The peroxisome-targeted PEX3Miro1 construct is sufficient for the mislocalization of components of the motor-adaptor complex to peroxisomes. This strategy permitted us to compare the recruitment of motor-adaptor complex components to peroxisomes in the presence of mutations of both Miro1 GTPase domains. Because endogenous wildtype Miro was not detectable on these peroxisomes, their distribution in the cells was governed entirely by the expressed mutations. This system also allowed us to compare functional differences between Miro1 and Miro2, as well as TRAK1 and TRAK2, in a manner not possible on mitochondria.

## Results

### Mislocalization of Miro1 is sufficient for co-localization of the motor-adaptor complex to peroxisomes

To redirect Miro1 to peroxisomes, we used a construct in which the C-terminal transmembrane domain of Miro1 had been removed and the transmembrane domain of peroxisome biogenesis factor, PEX3, had been added at the N-terminal of the construct (Basu et al., 2021). The N-terminal placement of the PEX3 transmembrane domain matches the orientation of the domain in PEX3 and localizes Miro1 to the peroxisome surface. Between the N-terminus of Miro1 and the PEX3 transmembrane domain, the construct contained a 278 amino acid segment consisting of an mRFP tag, a 6xhis tag, and an amino acid linker to create PEX3TM-6xhis-mRFP-Miro1 (hereafter called PEX3Miro1). We also made a control construct lacking Miro1 (PEX3TM-6xhis-mRFP, hereafter called PEX3-C).

Both PEX3Miro1 and PEX3-C were expressed in COS-7 cells with a peroxisome marker, mTurquoise-Serine-Arginine-Leucine (SRL), and appropriately localized to the outer membrane of peroxisomes (Figures 1A and S1 A,B). When expressed at high levels, however, both constructs were also present on mitochondria, a consequence of the PEX3 transmembrane domain and not the Miro1 sequence (Figure S1B); mitochondrial localization upon overexpression can occur with full-length PEX3 as well (Sugiura et al., 2017). In subsequent experiments, we therefore restricted our analysis to signals that co-localized with the mTurqoise-SRL marker (unless otherwise noted) to avoid any confounding effects of spillover onto mitochondria (see Methods) In addition, to minimize variability in our observations due to differences in PEX3-C and PEX3Miro1 expression across cells and bio-replicates, transfection and imaging conditions were defined with an initial set of samples and kept constant throughout. Under these defined conditions, cells having detectable amounts of PEX3Miro1 or PEX3-C co-localized with peroxisomes were considered for analysis.

**Figure 1.**
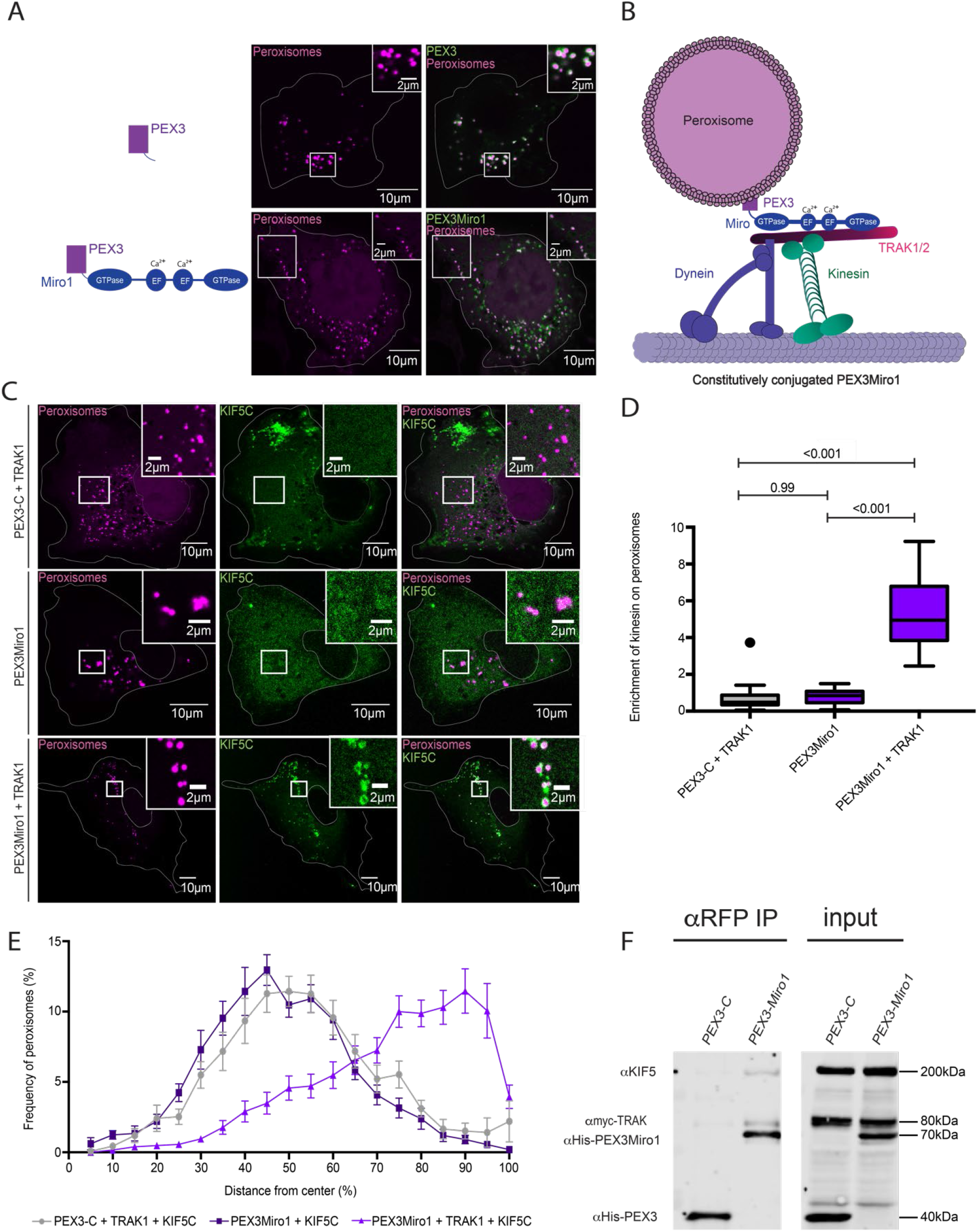
PEX3Miro1 localizes to peroxisomes and, with TRAK1 overexpression, can localize KIF5C to peroxisomes. A) In COS-7 cells, expressed PEX3-6xhis-mRFP (PEX3-C) or PEX3-6xhis-mRFP-Miro1 (PEXMiro1) (green) colocalizes with the mTurquoise-SRL peroxisome marker (magenta). B) Schematic of the PEX3Miro1 construct recruiting TRAK1/2 and KIF5C to the surface of a peroxisome. C) Expression of mTurquoise-SRL (magenta) with either PEX3-C or PEX3Miro1 and mCitrine-KIF5C (green) with or without co-expression of myc-TRAK1 in COS-7 cells. The presence of PEX3-C and PEXMiro1 on the peroxisomes was confirmed in each cell by imaging their RFP tags. D) Quantification of the amount of KIF5C enriched on peroxisomes that carry PEX3-C control or PEX3Miro1 with or without overexpression of myc-TRAK1. The quantifications here and subsequently are represented as ‘Box and Whiskers’ plots with the median value indicated. Outliers are represented as individual dots and are considered in statistical calculations. Here and in subsequent figures, P values are indicated and were determined by one-way ANOVA with Dunnett’s T3 correction for multiple comparisons. E) Distribution of peroxisomes marked with mTurquoise-SRL in the COS-7 cells analyzed in (D). Distance from the center of the cell in concentric shells was quantified using the DoveSonoPro FIJI macro (linked in methods). N=15 cells over 3 biological replicates. Error bars represent standard deviation. F) PEX3-C or PEX3Miro1 were expressed in HEK293T cells along with myc-TRAK1 and mCitrine-KIF5C and co-immunoprecipitated using antibodies to the RFP tag on the PEX3-C and PEX3Miro1 constructs. Western blots were stained using anti-His, myc, and KIF5 antibodies. Data from >3 biological replicates throughout. N=15 for panels A,C,D,E. The corner inserts show enlargement of the boxed regions.

To test whether the PEX3Miro1 construct was sufficient for relocalization of the motor-adaptor components to peroxisomes in COS-7 cells, we expressed either PEX3-C or PEX3Miro1 with mTurquoise-SRL and combinations of myc-TRAK1 and, the neuron-specific isoform of KIF5, KIF5C, that was tagged with mCitrine and YFP (hereafter called mCitrine-KIF5C) (Figure 1B,C,D). After fixation, the extent of mCitrine-KIF5C colocalization with mTurquoise-SRL was quantified using a custom FIJI macro (see Methods). KIF5C was diffuse in the cytosol or present on mitochondria but absent from peroxisomes when co-expressed with PEX3-C and myc-TRAK1. Both in Figure 1C and subsequent Figures, whether mCitrine-KIF5C was diffuse in the cytosol or also detectable on mitochondria varied from cell to cell and was dependent on expression level. In contrast, KIF5C was clearly present on peroxisomes when PEX3Miro1 and myc-TRAK1 were co-expressed (Figure 1C,D). Although variants of Miro1 have been reported to localize to peroxisomes, there was a clear lack of detectable mCitrine-KIF5C on peroxisomes in the absence of PEX3Miro1 (Costello et al., 2017; Covill-Cooke et al., 2020; Okumoto et al., 2018). This lack of peroxisomal mCitrine-KIF5C indicates that, if a peroxisome-targeted variant of Miro1 is endogenously present in these cells, it is not a significant factor and any KIF5C on peroxisomes is attributable to the PEX3Miro1 construct. These findings, and the ability to restrict our analysis to what is present on peroxisomes rather than on mitochondria, enable us to use the PEX3Miro system to study the regulation of the motor-adaptor complex without concern for the influence of endogenous mitochondrial Miro. When myc-TRAK1 was omitted, PEX3Miro1 was not sufficient to recruit KIF5C to peroxisomes (Figure 1C,D). We do not know why endogenous TRAK1 was not sufficient to permit PEX3Miro1 to recruit KIF5C; it may be that endogenous mitochondrial Miro1 outcompetes the peroxisomal transgene, but the finding confirms the requirement of TRAK1 in the motor-adaptor complex (Glater et al., 2006; Henrichs et al., 2020; Stowers et al., 2002; van Spronsen et al., 2013).

Peroxisomes in COS-7 cells typically reside near the nucleus. Although capable of microtubule-based transport (Huber et al., 1999; Rapp et al., 1996; Schrader et al., 1996; Wiemer et al., 1997), peroxisomes in COS-7 cells moved little and were seldom encountered in the periphery of a cell. This perinuclear localization of peroxisomes was evident in transfections where the expressed mCitrine-KIF5C was not on peroxisomes (PEX3-C and TRAK1 or PEX3Miro1 without TRAK1). In contrast, when we expressed PEX3Miro1 with myc-TRAK1 and mCitrine-KIF5C, peroxisomes moved more (as in (Basu et al., 2021)) and were more widely dispersed in the cell (Figure 1E). We quantified their distribution using a lab-developed image-analysis FIJI macro, DoveSonoPro (Basu et al., 2021). DoveSonoPro measures the % frequency of objects as they appear in concentric shells moving out from the center of the cell. The macro takes into consideration the shape of the cell and uses this shape to draw the concentric shells. Both the co-localization of KIF5C to peroxisomes and the increased peroxisome distribution were dependent on the co-expression of myc-TRAK1 with KIF5C (Figure 1E). As a further test of the ability of PEX3Miro1 to assemble the complex on peroxisomes, we expressed PEX3-C or PEX3Miro1 in HEK293T cells along with mCitrine-KIF5C and myc-TRAK1. myc-TRAK1 and mCitrine-KIF5C co-immunoprecipitated with PEX3Miro1 but not with PEX3-C control (Figure 1F). These results establish that PEX3Miro1 is sufficient for localizing the motor-adaptor complex and that the complex is active, i.e. capable of moving peroxisomes in the direction of microtubule + ends, as expected for the KIF5C motor.

### The N-terminal GTPase determines Miro1 interactions with the mitochondrial motor-adaptor complex

The localization of a functional motor-adaptor complex to peroxisomes by PEX3Miro1 allowed us to examine the significance of GTPase mutations without confounding effects of endogenous Miro. We introduced single point mutations to the GTPase domains of the PEX3Miro1 construct to confer either a constitutively active GTP-state (N-GTPase:P13V or C-GTPase:K427N) or a constitutively inactive GDP-state (N-GTPase:T18N or C-GTPase:S432N) (Fransson et al., 2006) (Figure 2). The constitutively active mutation locks the N-GTPase in a GTP-bound conformation, as confirmed by crystallography (Smith et al., 2020). The inactive N-GTPase mutant locks the GTPase in a predicted GDP-bound conformation where GTP should no longer be able to bind.

**Figure 2.**
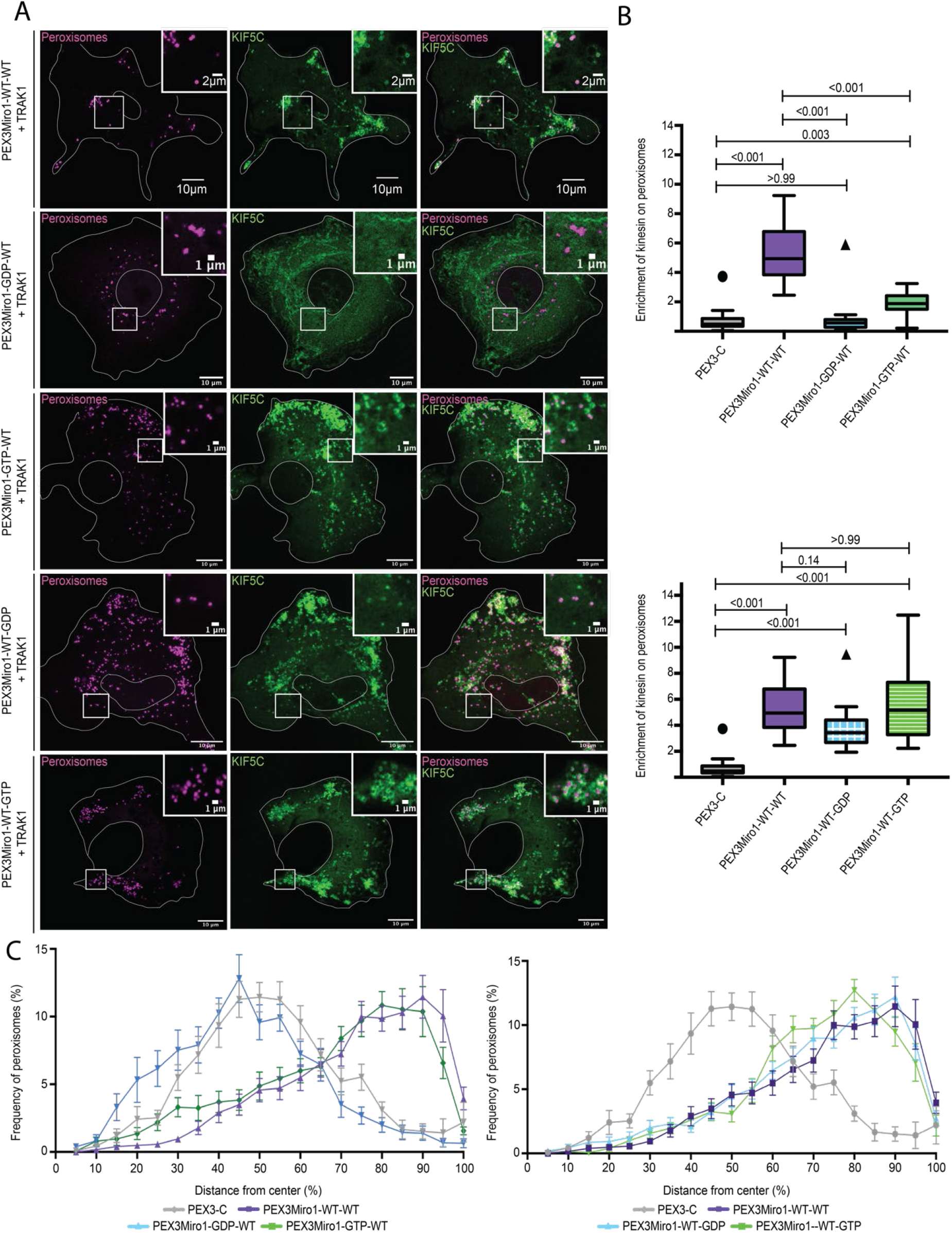
The GTPase domains of Miro1 regulate co-localization with KIF5C on peroxisomes. A) Expression in COS-7 cells of mTurquoise-SRL (magenta), myc-TRAK1, and mCitrine-KIF5C (green) with either PEX3Miro1wild-type or PEX3Miro1 carrying a mutation of either the N-or C-GTPase domains: N- GDP-state T18N, N- GTP-state P13V, C-GDP-state S432N, C- GTP-state K427N. The presence of the PEXMiro1constructs on peroxisomes was confirmed in each cell by imaging their RFP tags. The corner inserts show enlargement of the boxed regions. Here and in subsequent figures the status of the two GTPase domains is abbreviated as either WT, GTP, or GDP for first the N-domain and then the C-domain. B) Quantification of the amount of KIF5C enriched on peroxisomes carrying PEX3-C and the constructs shown in (A) For clarity, the mutations of the N-GTPase domain (above) are plotted separate from those of the C-GTPase (below) though transfected and imaged in the same experiments with the same controls. The quantification is represented as ‘Box and Whiskers’ plots with the median value indicated. Outliers are represented as single plot points and were included in all statistical calculations. Here, and in all subsequent figures, P values were determined by one-way ANOVA with Dunett’s T3 correction for multiple comparisons. N=15 cells over 3 biological replicates. C) Quantification of peroxisomal distribution in concentric shells radiating from the center of the cell in the same cells analyzed in (B), with separate plots for mutations of the N-GTPase (left) and C- GTPase (right) for clarity, as in (B). N=15 cells over 3 biological replicates. Error bars represent standard deviation. The data for the negative control (PEX3C) and positive control (PEX3Miro1WT-WT) are repeated from Figure 1 for clarity and are from experiments conducted at the same time.

To determine whether the N-GTPase domain regulates the assembly of the motor-adaptor complex, we quantified the effect of the PEX3Miro1 mutations on the localization of mCitrine-KIF5C to peroxisomes. The mRFP tag on these constructs confirmed that the mutated Miro constructs still localized to peroxisomes and thus a failure to localize KIF5C to peroxisomes would reflect a failure of the complex to assemble on PEX3Miro1. When the N-GTPase carried the GDP-state T18N mutation, PEX3Miro1 no longer caused KIF5C to go to peroxisomes when TRAK1 was co-expressed, whereas the GTP-state mutation (P13V) did (Figure 2A, B). Quantitatively, however, the GTP-state mutant did not recruit as much KIF5C as wild-type PEX3Miro1 (Figure 2B). This widely used constitutively active GTP mutation thus may not be perfectly equivalent to wild-type PEX3Miro1 with a bound GTP; in addition to blocking the GTPase activity, it may slightly alter the structure of the domain and diminish its function. The cellular distribution of peroxisomes in these experiments was altered by successful KIF5C recruitment. Peroxisomes moved to the periphery in cells expressing wild-type or the GTP-state mutant of the N-GTPase of PEX3Miro1 but not in cells expressing the GDP-state mutant or the PEX3-C control (Figure 2D).

### Miro1 C-terminal GTPase regulation of the mitochondrial motor-adaptor complex

Miro1 C-GTPase mutants have previously been overexpressed and altered mitochondrial distribution through an unknown mechanism (Fransson et al., 2006). We expressed PEX3Miro1 C-GTPase mutants in COS-7 cells together with mTurquoise-SRL, myc-TRAK1, and mCitrine-KIF5C and quantified KIF5C on peroxisomes. The mRFP tag on PEX3Miro1 confirmed that it was expressed on peroxisomes. KIF5C was also present on peroxisomes in each of the C-GTPase mutants (Figure 2A,C). There were, however, possibly subtle effects on peroxisomal KIF5C in the S432N C-GTPase GDP-state mutant (Figure 2C) relative to wildtype or the K427N GTP-state mutant of PEX3Miro1 imaged in the same experiment. Consistent with the very small effects of the C-GTPase mutations on KIF5C recruitment, they did not cause differences in the distribution of peroxisomes bearing them (Figure 2E).

To further investigate a possible role for the PEX3Miro1 C-GTPase, we made PEX3Miro1 double mutants with both the N-GTPase and C-GTPase in either the GTP-state or GDP-state. Upon co-expression in COS-7 cells with mTurquoise-SRL, myc-TRAK1, and mCitrine-KIF5C, each double mutant matched the corresponding mutations of the N-GTPase alone. Peroxisomes with both domains in the GDP-state did not co-localize with mCitrine-KIF5C but those with both domains in the GTP-state did (Figure 3A,C). We also made double mutants in which the two domains were locked in opposite states (Figure 3B,C). Again, the N-GTPase was the primary determinant of KIF5C recruitment; all the constructs in which the N-GTPase was in the GTP-state recruited KIF5C to peroxisomes and shifted their distribution to the periphery and all those in which the N-GTPase was in the GDP-state did not (Figure 3B-D). Quantification, however, found that the construct with N-GTPase in the GTP-state was slightly more effective in recruiting KIF5C to peroxisomes when the C-GTPase was in the GTP-state than when it was in the WT or GDP-state, (Figure 3C). Together, these results indicate that the state of PEX3Miro1 N-GTPase very largely determines the ability to recruit KIF5C.

**Figure 3.**
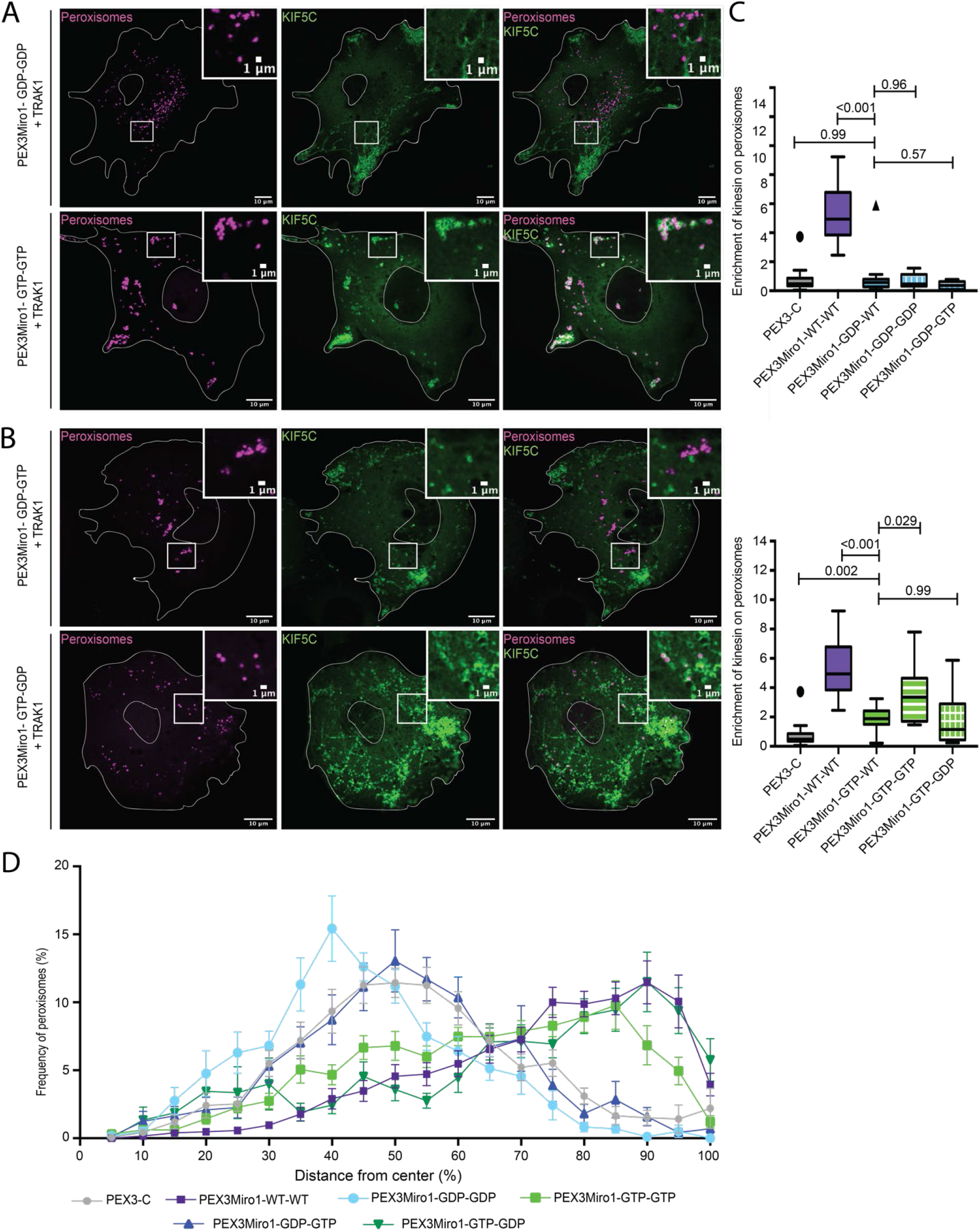
The N-GTPase of Miro1 has a predominant influence on KIF5C recruitment. A) Expression of PEX3Miro1 N-GTPase and C-GTPase double mutants in which both domains were in either the GTP or GDP state and its consequences for mCitrine-KIF5C (green) recruitment to peroxisomes (mTurquoise-SRL, magenta) with co-expression of myc-TRAK1. The corner inserts show enlargement of the boxed regions. B) Expression PEX3Miro1 N-GTPase and C-GTPase double mutants in which the two domains were in opposite states and its consequences for mCitrine-KIF5C (green) recruitment to peroxisomes (mTurquoise-SRL, magenta) with co-expression of myc-TRAK1. In (A, B) the presence of the PEXMiro1 constructs on peroxisomes was confirmed in each cell by imaging their RFP tags. C) Quantification of the amount of KIF5C enriched on cells transfected as in (A) and (B). If the N- GTPase is in the GDP-bound state (upper graph) or in the GTP-state (lower graph), the state of the C-GTPase has little or no effect, except for a modest enhancement of KIF5C recruitment when both domains are GTP-bound rather than only the N-GTPase. The quantification is represented as ‘Box and Whiskers’ plots with the median value indicated. Outliers are represented as single plot points and were included in all statistical calculations. Indicated P values are determined by one-way ANOVA with Dunnett’s T3 correction for multiple comparisons. N=15 cells over 3 biological replicates. D) Quantification of peroxisome distribution from the cells in (A-D). Error bars represent standard deviation. The data for the negative control (PEX3C) and positive control (PEX3Miro1WT-WT) are repeated from Figure 1 for clarity and are from experiments conducted at the same time.

### The Miro1 N-GTPase domain regulates recruitment of P135

To examine the influence of the GTPase domains on the ability of the motor adaptor complex to interact with the dynein/dynactin retrograde motor, we assayed the distribution of P150^Glued^, a microtubule-binding protein that is the largest subunit of dynactin. mCitrine-tagged P150^Glued^ however, when expressed in COS7 cells, coats all the microtubules, which made it impossible to determine if it was also present on PEX3Miro1-expressing peroxisomes. We therefore used mCitrine-tagged P135, a construct that lacks the microtubule-binding motif (Dixit et al., 2008; Tokito et al., 1996), and quantified P135 on peroxisomes tagged with mTurquoise-SRL. Like the KIF5C experiments, we found that PEX3Miro1 but not PEX3-C could recruit P135 to peroxisomes and that overexpression of TRAK1 was necessary for this recruitment (Figure 4A,B). P135 recruitment was also assayed with the GTPase mutations used in figures 2 and 3. When the PEX3Miro1 N-GTPase was wild-type or carried the GTP-state mutation, P135 was recruited to peroxisomes. The N-GTPase GDP-state mutation, however, did not recruit P135 (Figure 4A,C). Mutation of the C-GTPase to either the GTP or GDP state had less effect on P135 (Figure 4A,D). Thus, the regulation of P135 recruitment followed the same rules as KIF5C and was predominantly dependent on the state of the N-GTPase domain.

**Figure 4.**
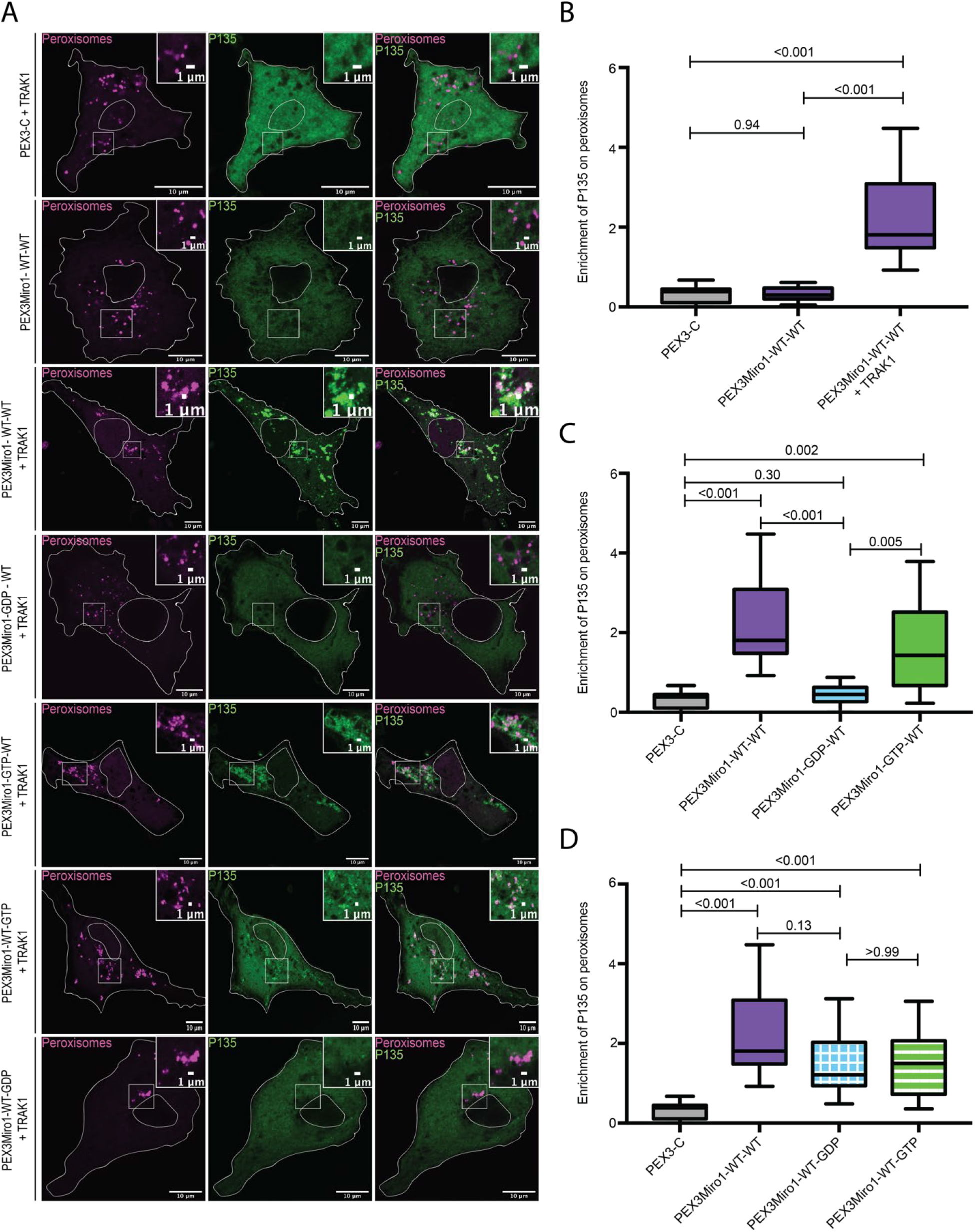
The GTPase domains of Miro1 regulate co-localization with P135. A) PEX3-C, PEX3Miro1, or PEX3Miro1 with mutations of either the N- or C- GTPase domains as indicated, and with or without TRAK as indicated were expressed in COS-7 cells with mCitrine-P135(green) and the peroxisome marker mTurquoise-SRL (magenta). The presence of the PEXMiro1 constructs on peroxisomes was confirmed in each cell by imaging their RFP tags. The corner inserts show enlargement of the boxed regions. (B-D) P135 colocalization with peroxisomes quantified from cells transfected as in (A) and represented as ‘Box and Whiskers’ plots with the median value indicated. Outliers are plotted as single points and were included in all statistical calculations. The indicated P values were determined by one-way ANOVA with Dunnett’s T3 correction for multiple comparisons. N=15 cells over 3 biological replicates. The data for the negative control (PEX3C) and positive control (PEX3Miro1WT-WT) are repeated in each graph for clarity and are from experiments conducted at the same time. B) Quantification of the amount of P135 enriched on PEX3-C control or PEX3Miro1 peroxisomes with or without expression of myc-TRAK1. C) Quantification of the amount of P135 enriched on peroxisomes bearing PEX3Miro1 wild-type or the PEX3Miro1 N-GTPase mutants. PEX3Miro1 constructs were co-expressed with myc-TRAK1 and mCitrine-P135. D) Quantification of the amount of KIF5C enriched on peroxisomes bearing PEX3Miro1 wild-type or the PEX3Miro1 C-GTPase mutants. PEX3Miro1 constructs were co-expressed with myc-TRAK1 and mCitrine-P135.

### The Miro1 N-GTPase regulates co-localization of TRAK1 and TRAK2 with PEX3Miro1

Consistent with previous models of the motor/adaptor complex, our experiments indicated that TRAK was required for the association of the motors with Miro. We therefore hypothesized that the GDP-state of the N-GTPase prevented TRAK association with Miro and the absence of TRAK on the peroxisomes accounted for the failure to recruit KIF5C and P135. We co-expressed in COS7 cells the PEX3Miro1 N-GTPase mutants with mTurquoise-SRL and TRAK1 that was tagged with mCitrine and YFP (hereafter called mCitrine-TRAK1) and quantified the amount of mCitrine-TRAK1 on peroxisomes. Consistent with our hypothesis, the PEX3Miro1 N-GTPase wild-type and GTP-state mutant were both able to recruit TRAK1 to peroxisomes whereas the N-GTPase GDP-state mutant could not (figure 5A,B).

**Figure 5.**
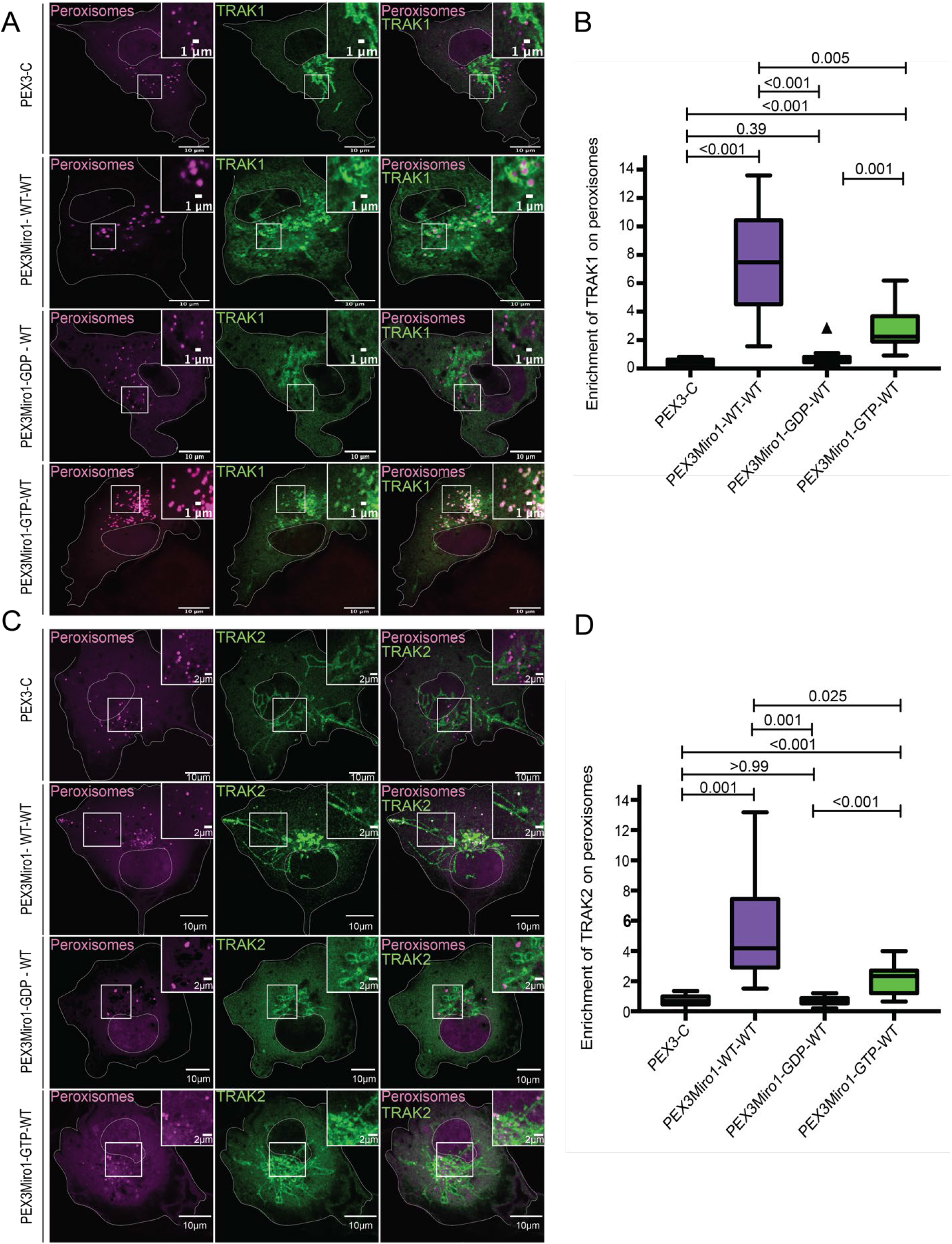
The PEX3Miro1 N-GTPase regulates co-localization of TRAK1 and TRAK2 with PEX3Miro1. A) PEX3-C, PEX3Miro1, or PEX3Miro1 with N-GTPase GDP and GTP-state mutants were expressed in COS-7 cells with mCitrine-TRAK1 (green) and mTurquoise-SRL (magenta). The presence of the PEXMiro1 and PEX3-C constructs on peroxisomes was confirmed in each cell by imaging their RFP tags. B) From cells as in (A), quantification of the amount of TRAK1 enriched on peroxisomes bearing PEX3-C, PEX3Miro1 and PEX3Miro1 N-GTPase mutants. C) PEX3-C, PEX3Miro1, or PEX3Miro1 with N-GTPase GDP and GTP-state mutants were expressed in COS-7 cells with mCitrine-TRAK2 (green) and mTurquoise-SRL (magenta). The presence of the PEXMiro1 constructs on peroxisomes was confirmed in each cell by imaging their RFP tags. D) From cells as in (C), quantification of the amount of TRAK2 enriched on peroxisomes bearing PEX3-C, PEX3Miro1, and PEX3Miro1 N-GTPase mutants. In (B) and (D) TRAK colocalization with peroxisomes is represented as ‘Box and Whiskers’ plots with the median value indicated. Outliers are plotted as single points and were included in all statistical calculations. The indicated P values are from analysis with One-way ANOVA. N=15 cells over 3 biological replicates. N=15 for all panels. The corner inserts show enlargement of the boxed regions.

Previous studies have indicated that the two isoforms of TRAK, TRAK1 and TRAK2, have functional differences (Brickley et al., 2005; Macaskill et al., 2009b; Quintanilla et al., 2020; van Spronsen et al., 2013). The PEXMiro assay allowed us to ask whether TRAK2 would also be recruited to PEXMiro1-expressing peroxisomes and subject to the same regulation by the GTPase domains. We therefore co-expressed mCitrine-TRAK2 with PEX3Miro1 and mTurquoise-SRL. Like TRAK1, TRAK2 localized to peroxisomes with wildtype PEXMiro1 and PEXMiro1 with the N-GTPase in the GTP-state, but not in the GDP-state (Figure 5C,D). The two isoforms also behaved similarly when the C-GTPase and double GTPase mutants were tested. Quantitatively, mCitrine-TRAK1 and mCitrine-TRAK2 on peroxisomes responded to the state of the Miro1 GTPase domains in the same way as mCitrine-KIF5C (Figure S2,S3). We conclude that the N-GTPase of Miro1 controlled motor recruitment by determining whether or not Miro1 bound a TRAK.

### TRAK1 and TRAK2 differ in their ability to recruit KIF5C to peroxisomes

The requirement for overexpressed TRAK as an adaptor for the recruitment of KIF5C in the peroxisome system allowed us to directly compare the function of TRAK1 and TRAK2 in serving as adaptors for KIF5C. We co-expressed PEX3Miro1 with mTurquoise-SRL, mCitrine-KIF5C, and either myc- TRAK1 or myc- TRAK2. TRAK2 expression could also localize KIF5C to peroxisomes (Figure S4), but comparatively TRAK2 recruited significantly less KIF5C than TRAK1 (Figure 6A,B). This difference was not due to different amounts of TRAK1 and 2 on the peroxisomes; co-expression of either mCitrine-TRAK1 or mCitrine-TRAK2 with PEX3Miro1 resulted in similar amounts of TRAK on peroxisomes (Figure 6C,D). We also quantified the ratio of peroxisomal KIF5C to peroxisomal TRAK1 and TRAK2 by co-expressing PEX3Miro1 with an mTurqouise-KIF5C construct in the presence of either mCitrine-TRAK1 or mCitrine-TRAK2. We normalized the amount of KIF5C to the amount of each TRAK and again found that more KIF5C co-localized with TRAK1 than TRAK2 (Figure S5). Thus, in comparing the two TRAK isoforms, the difference in KIF5C recruitment to peroxisomes is dependent on differences in the TRAK-KIF5C interactions and not differences in TRAK expression or TRAK-Miro interactions.

**Figure 6.**
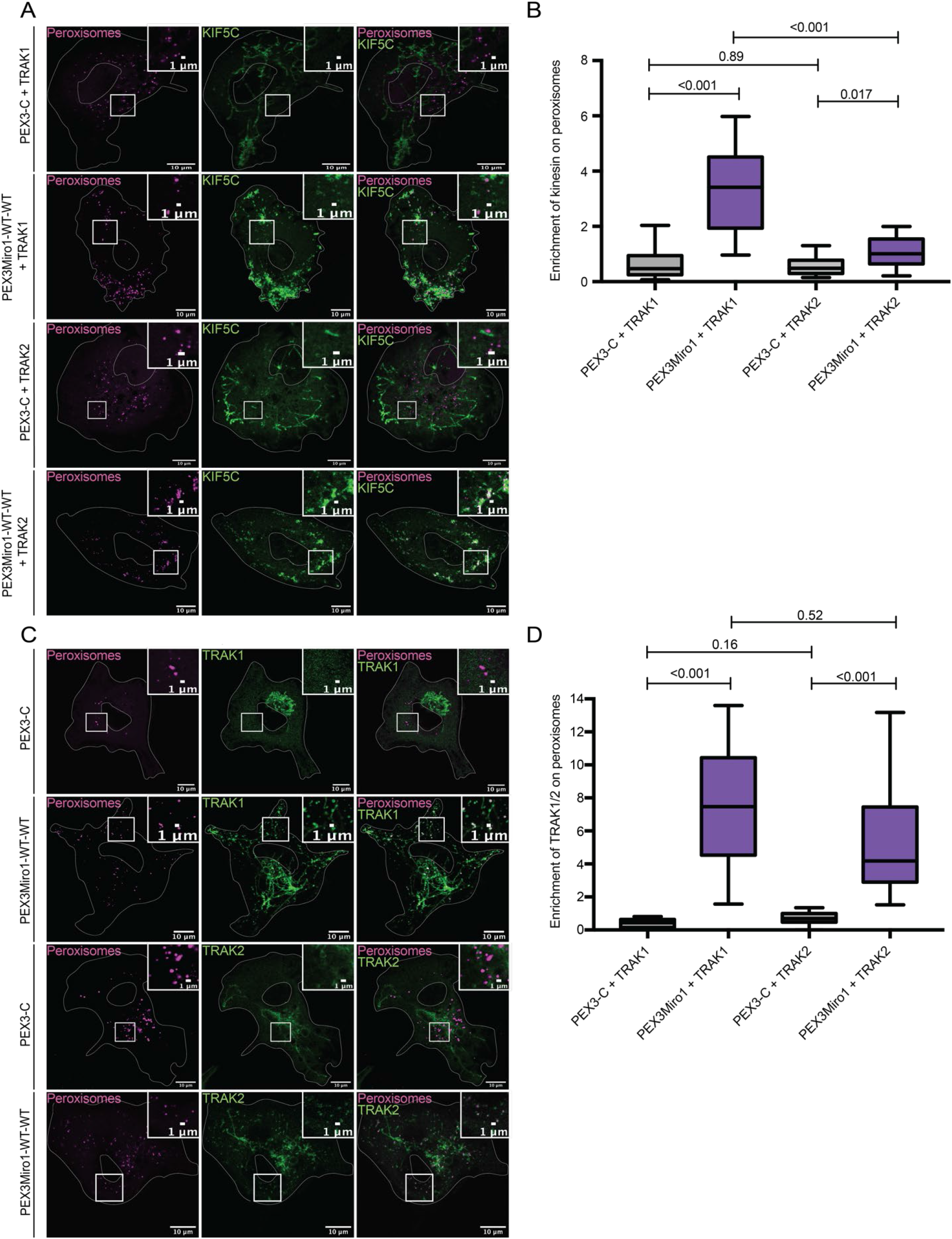
TRAK1 and TRAK2 both serve as adaptors for KIF5C but differ in their ability to recruit KIF5C. A) Either myc-TRAK1 or myc-TRAK2 were expressed in COS-7 cells together with mCitrine-KIF5C (green), mTurquoise-SRL (magenta) and either PEX3-C or PEX3Miro1. The presence of the PEXMiro1 or PEX3-C constructs on peroxisomes was confirmed in each cell by imaging their RFP tags. The corner inserts show enlargement of the boxed regions. B) From cells transfected as in (A), quantification of the amount of KIF5C enriched on peroxisomes with either mCitrine-TRAK1 or mCitrine-TRAK2 was expressed in COS-7 cells together with mTurquoise-SRL (magenta) and either PEX3-C or PEX3Miro1. The presence of PEXMiro1 or PEX3-C on peroxisomes was confirmed in each cell by imaging their RFP tags. C) Either mCitrine-TRAK1 or mCitrine-TRAK2 (green) were expressed in COS-7 cells together with mTurquoise-SRL (magenta) and either PEX3-C or PEX3Miro1. The presence of the PEX3Miro1, PEX3-C constructs on peroxisomes was confirmed in each cell by imaging their RFP tags. D) From cells transfected as in (C), quantification of the amount of mCitrine-TRAK1 or mCitrine-TRAK2 enriched on peroxisomes. In (B) and (D), colocalization with peroxisomes is represented as ‘Box and Whiskers’ plots with the median value indicated. The indicated P values were determined by one-way ANOVA with Dunnett’s T3 correction for multiple comparisons. N=15 cells over 3 biological replicates.

### PEX3Miro2 does not recruit the motor-adaptor complex to peroxisomes

In mammals, Miro1 and Miro2 are 60% identical (Fransson et al., 2003; Fransson et al., 2006). The extent to which their functions differ remains unclear and the role of Miro2 in mitochondrial motility has not received as much attention. The PEX3Miro approach afforded an opportunity to compare Miro1 and Miro2. We created a PEX3TM-6xhis-mRFP-Miro2 construct (PEX3Miro2) equivalent to PEX3Miro1 to ask whether this peroxisomal Miro2 would interact with TRAK and kinesin. We expressed PEX3-C, PEX3Miro1, or PEX3Miro2 together with mTurquoise-SRL and mCitrine-KIF5C and with or without myc-TRAK1. PEX3Miro2, however, was not able to recruit KIF5C to peroxisomes whether or not myc-TRAK1 was co-expressed, although PEX3Miro1 with myc-TRAK1, assayed in parallel, recruited KIF5C as expected (Figure 7A). PEX3Miro2 also failed to recruit mCitrine-TRAK1 or mCitrine-TRAK2 to peroxisomes (Figure S6). To determine whether the reason that PEX3Miro2 behaved so differently from PEX3Miro1 was due to the state of the GTPase domains it assumed when expressed in COS7 cells, we introduced mutations into the N- and C-GTPase domains of PEX3Miro2 that would create the GTP and GDP states for each domain. These constructs were co-expressed with mTurquoise-SRL and myc-TRAK1 and mCitrine-KIF5C. None of the GTPase mutations in PEX3Miro2 enabled the recruitment of the motor-adaptor complex to peroxisomes (Figure 7A,B). The unexpected difference in the behavior of Miro1 and Miro2 in this assay is thus unrelated to either the choice of TRAK isoform as a partner nor the state of the GTPase domains.

**Figure 7.**
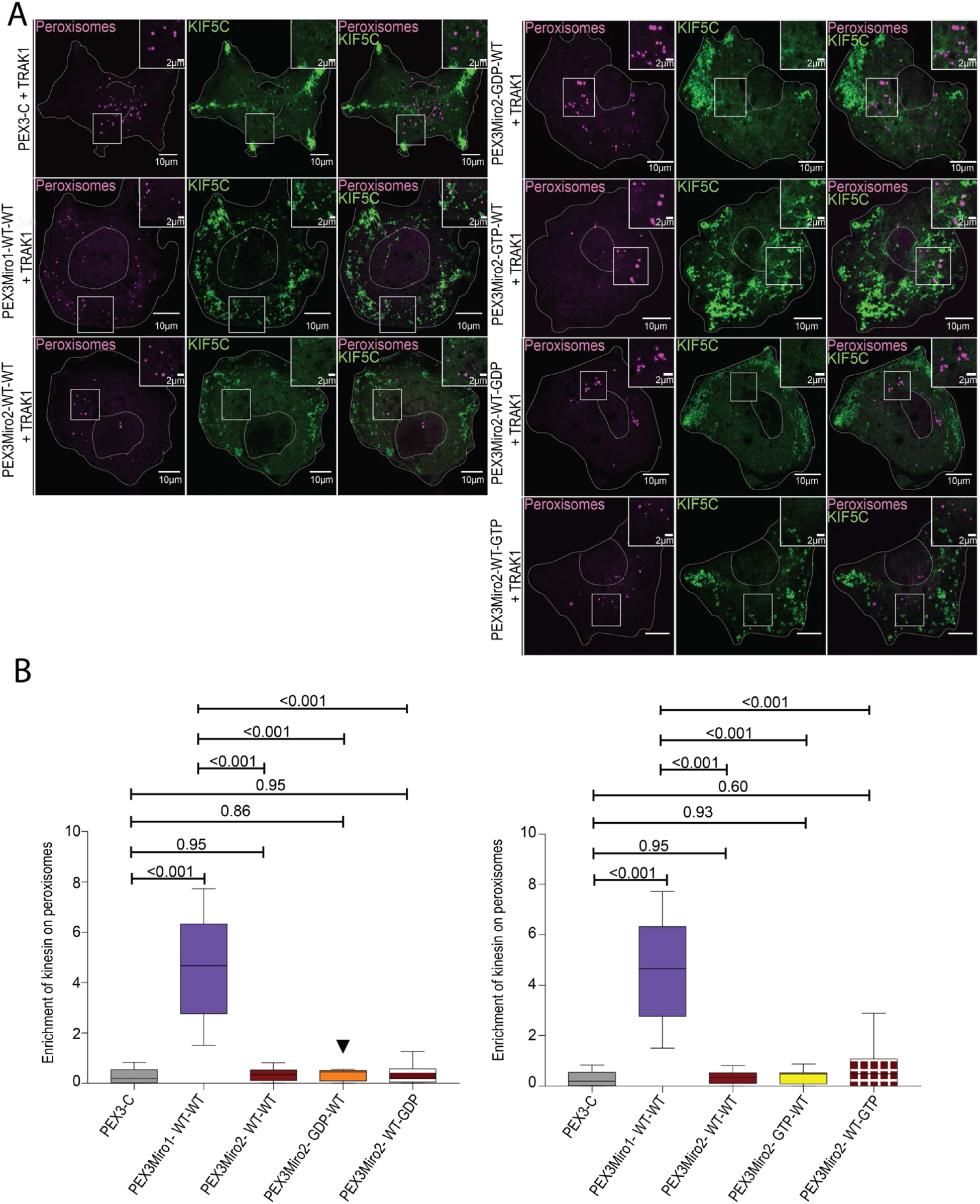
PEX3Miro2 does not recruit KIF5C to peroxisomes. A) Expression in COS-7 cells of mTurquoise-SRL (magenta), myc-TRAK1, and mCitrine-KIF5C (green) with either PEX3-C, PEX3Miro1, PEX3Miro2, or PEX3Miro2 carrying a mutation of either the N- or C- GTPase domains: N- GDP-state T18N, N- GTP-state A13V, C-GDP-state S430N, C- GTP-state A425V. The presence of the PEX3Miro constructs on peroxisomes was confirmed in each cell by imaging their RFP tags. The corner inserts show enlargement of the boxed regions. B) Quantification of the amount of KIF5C enriched on peroxisomes carrying the constructs shown in (A). For clarity, the mutations to GDP-state and GTP-state are plotted separately. The quantification is represented as ‘Box and Whiskers’ plots with the median value indicated. Outliers have been removed from this data set using the ROUT method (Q=1%) and are not included in statistical calculations. Indicated P values were determined by one-way ANOVA with Dunnett’s T3 correction for multiple comparisons. N=15 cells over 3 biological replicates.

### PEX3Miro1 can cause the redistribution of peroxisomes in hippocampal neurons

The behavior of PEXMiro1 in COS-7 cells suggested that it might also be able to alter the motility and distribution of peroxisomes in neurons and that this effect would depend on the state of the N-GTPase domain. We therefore co-expressed PEX3Miro1 and PEX3Miro1 N-GTPase mutants along with mTurquoise-SRL, myc-TRAK1, and mCitrine-KIF5C in rat hippocampal neurons. Neurons were transfected on DIV3 and fixed on DIV5. To quantify their distribution, we counted the number of peroxisomes in the soma and in distal neurites. In control neurons expressing PEX3-C peroxisomes are present throughout the neuron but are mostly localized to the soma. When we express wild-type PEX3Miro1 along with myc-TRAK1 and mCitrine-KIF5C, the peroxisomes undergo a substantial re-distribution, almost entirely leaving the soma and accumulating instead in distal axons and at growth cones (Figure 8, Figure S7). This re-distribution is consistent with the KIF5C-mediated movement of the peroxisomes to the + ends of the axonal microtubules. The same redistribution occurs with the N-GTPase GTP-state mutant. In contrast, upon expression of the N-GTPase GDP state mutant with myc-TRAK1 and mCitrine-KIF5C, the vast majority of the peroxisomes accumulated in the soma and very few were in the neurites (Figure 8). Thus, the Miro1 N-GTPase can alter organelle distribution in hippocampal neurons and is likely to be a major determinant of mitochondrial behavior in both neurons and non-neuronal cells.

**Figure 8:**
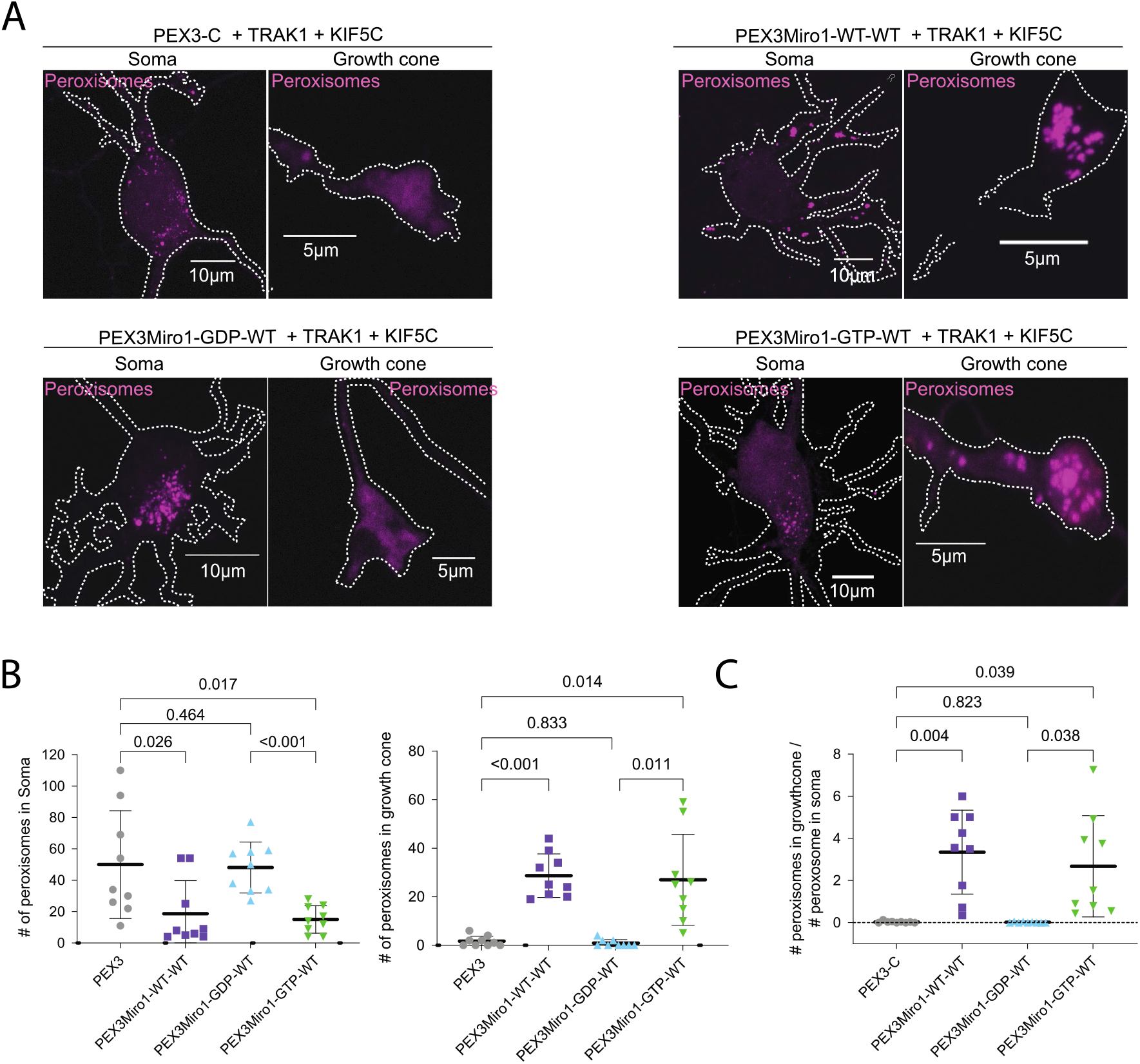
PEX3Miro1-dependent redistribution of peroxisomes in hippocampal neurons is regulated by the N-GTPase. A) Expression of mTurquoise-SRL peroxisome marker (shown in magenta) with either PEX3 control, wild-type PEX3Miro1, or PEX3Miro1 nGTPase mutants together with myc-TRAK1 (signal not shown) and mCitrine-KIF5C (signal not shown) in soma and growth cone of rat hippocampal neurons. Cells were fixed at DIV5. B) Quantification of the distribution of peroxisomes from neurons as in (A). For each neuron, peroxisomes within a single representative growth cone or soma were counted. All data points are plotted, N=9. For each dataset, line indicates mean and whiskers indicates SD. The indicated P values were obtained one-way ANOVA with Dunnett’s T3 correction for multiple comparisons. C) Quantification of the distribution of peroxisomes from neurons as in (A). For each neuron, peroxisomes within a single representative growth cone were counted and then expressed as a ratio relative to the number of peroxisomes in the soma of that neuron. All data points are plotted, N=9. The indicated P values were obtained by one-way ANOVA with Dunnett’s T3 correction for multiple comparisons.

## Discussion

By misdirecting Miro to the surface of peroxisomes, this study has allowed us to quantify the recruitment of other components of the motor-adaptor complex to Miro and to observe the consequences of that recruitment on peroxisomal distribution. The relocalization of TRAK and the motors to peroxisomes served as a readout of the ability of the complex to assemble and thereby permitted us to independently examine the two isoforms of Miro and of TRAK in a cellular milieu and reveal differences in their ability to support assembly of the complex. In addition, the system revealed that perturbations of the N-GTPase domain of Miro altered assembly of the complex. These findings revealed a mechanistic basis for the previously reported influences of overexpressed GTPase-domain mutations on mitochondrial distribution (Fransson et al., 2003; Fransson et al., 2006; MacAskill et al., 2009a). The N-GTPase domain is a crucial regulator of microtubule-based motility, much as Miro’s EF hands were previously shown to mediate the Ca^2+^-dependent arrest of that motility (Macaskill et al., 2009b; Saotome et al., 2008; Wang and Schwarz, 2009).

By re-directing Miro constructs to peroxisomes, we avoided altering mitochondrial health or motility; mitochondria retained their endogenous Miro proteins. Though our Miro constructs, particularly if expressed at high levels, could also be found on mitochondrial membranes despite the replacement of their mitochondrial transmembrane domains with a peroxisome-targeting sequence, by restricting our analysis to peroxisomes, we could study the influence of isoforms and mutations without the confounding presence of endogenous motor-adaptor proteins on mitochondria. The “spillover” of highly expressed PEX3Miro constructs onto mitochondria was not a consequence of the Miro domains; overexpressed PEX3 alone also resided on both mitochondria and peroxisomes and to the same extent as PEX3Miro (Figure S1). One drawback of this “spillover” onto mitochondria, however, was that the PEX3Miro system was not suitable for biochemical analysis of complex assembly because too much of the protein was present on mitochondria and likely co-assembled there with endogenous Miro. We therefore restricted our analysis to colocalization of components with the PEX3Miro-bearing peroxisomes where the ability to recruit either TRAK or a motor was strictly dependent on the nature of the PEX3Miro construct.

Both TRAK isoforms and KIF5C co-localized with peroxisomes that carried the RFP-tagged PEX3Miro1 construct and this was accompanied by a shift in the peroxisome localization to the periphery of the cell. We never saw either TRAK or KIF5C on peroxisomes when the PEX3 transmembrane domain without Miro1 (the PEX3-C control) was expressed, nor did PEX3-C cause the same redistribution of peroxisomes within the cell (Figure 1). Thus, although several variants of Miro1 have been reported to be expressed on peroxisomes (Costello et al., 2017; Covill-Cooke et al., 2020; Okumoto et al., 2018), if they were present on peroxisomes in our COS-7 cells they were at too low levels to influence our assays. The co-localization data and changes in distribution were due to the PEX3Miro1 construct and this was borne out by subsequent studies in which mutations of PEX3Miro1 prevented recruitment of both TRAKs and motors. Recently, a PEX26-Miro1 fusion was reported that similarly altered peroxisome distribution and, as in our experiments, it was found that the redistribution depended on the GTPase state of Miro1’s N-GTPase domain (Castro and Schrader, 2018). Our experiments provide a mechanistic explanation of this phenomenon by showing that motor-recruitment was strictly dependent on the presence of TRAK and that TRAK could not associate with Miro1 whose N-GTPase was locked in the GDP-binding state.

The expression of PEX3Miro1 sufficed to recruit co-expressed TRAK1 and KIF5C or P135 to peroxisomes. Recently it has been reported that two mitochondrial transmembrane proteins, Metaxins1 and 2, can also be involved in the assembly of the motor-adaptor complex (Zhao et al., 2021). We do not know whether the recruitment we observed was independent of Metaxins, peroxisomes contain endogenous Metaxins, or PEX3Miro1 also caused Metaxins to relocate to peroxisomes. If Metaxins are needed for correct insertion and organization of Miro on mitochondria, akin to their other known functions in the mitochondrial translocation apparatus (Abdul et al., 2000; Armstrong et al., 1997), it is possible that they are not needed when their membrane-association is driven by the PEX3 transmembrane domain. On the other hand, we note that PEX3Miro1 did not recruit sufficient endogenous TRAK to drive motor recruitment, which suggested that the endogenous mitochondrial Miro outcompeted PEX3Miro for TRAK. A higher affinity of mitochondrial Miro for TRAK might reflect an influence of Metaxins and further studies with PEX3Miro1 may clarify their role in the motor adaptor complex.

The function of Miro’s GTPase domains have been of interest since Miro’s discovery and was examined chiefly by overexpressing Miro1 GTP and GDP-state mutants. N-terminal GTPase mutants altered mitochondrial distribution and morphology in cell lines and neurons. (Fransson et al., 2003; Fransson et al., 2006; MacAskill et al., 2009a). In a Drosophila Miro loss-of-function mutant, the N-GTPase GDP-state mutant led to an accumulation of mitochondria in the soma of neurons and ultimately led to premature lethality (Babic et al., 2015). In Miro1/Miro2 double knockout MEFs, the Miro1 N-GTPase GTP-state mutant partially rescued mitochondrial motility whereas the GDP-state mutant did not (Norkett et al., 2020). We offer here a mechanism behind those observations: the GDP-bound state failed to interact with either TRAK1 or TRAK2 and TRAK was absolutely required for recruiting the motors to Miro1. The N-GTPase of Miro1 also regulates the interaction between Miro1 and DISC1, CENP-F, and Myo19 (Kanfer et al., 2015; Norkett et al., 2020; Oeding et al., 2018). In these cases, as with TRAK, the Miro1 N-GTPase GDP-state mutant prevented the interactions of Miro1 that occur with wild-type Miro1 and the GTP-state mutant. Further experiments with the PEX3Miro system may determine whether the interactions of Myo19, DISC1, and CENP-F with Miro are all facilitated by the presence of TRAK or whether their dependence on the GTP-state of the N-GTPase is a TRAK-independent phenomenon.

We have only very small differences between wild-type Miro1 and the Miro1 C-GTPase mutants’ ability to recruit kinesin to peroxisomes. The influence of the C-GTPase on complex formation was very minor compared to that of the N-GTPase; in isolation, neither C-GTPase mutant differed from wildtype PEX3Miro1 in their ability to recruit TRAK1 and KIF5C to peroxisomes and mutations of the C-GTPase could not rescue their recruitment when the N-GTPase was in the GDP state. However, when the N-GTPase carried the GTP state mutation P13V – a state that was less efficient at recruiting TRAK1 and KIF5C – the GTP state of the C-GTPase made some difference. Heretofore, there has been little evidence for a function of the C-GTPase. Mutation of this domain has not been shown to alter interactions with Myo19, DISC1, or CENP-F (Kanfer et al., 2015) and mutations of the Miro1 C-GTPase overexpressed in COS-7 cells showed no difference in mitochondrial distribution or cell health (Fransson et al., 2003; Fransson et al., 2006). An exception was reported in *Drosophila* where the C-GTPase GDP-state mutation decreased retrograde motility (Babic et al., 2015). More quantitative assays may detect further C-GTPase effects. From our experiments, we infer that the GTP-state of the C-GTPase may promote motor complex assembly, but that it is a subtle influence compared to the clear requirement of the GTP state for the N-GTPase.

The quantitative nature of our system also allowed us to examine differences between TRAK1 and TRAK2 while keeping the other components of the complex constant. The importance of the TRAKs and their *Drosophila* homolog Milton has been clear from loss of function phenotypes and protein interaction studies (Glater et al., 2006; MacAskill et al., 2009a; Randall et al., 2013; Stowers et al., 2002). Functional differences in the isoforms were first noted in *Drosophila* where one splicing variant, Milton-C, failed to co-immunoprecipitate with kinesin or recruit kinesin to mitochondria when expressed in COS-7 cells (Glater et al., 2006). Subsequent mammalian studies have reported functional differences between TRAK1 and 2, with TRAK1 seeming to be the primary adaptor for KIF5C driven plus-end directed trafficking (Brickley et al., 2005; MacAskill et al., 2009a; van Spronsen et al., 2013). The difference in kinesin binding arises from a TRAK2 conformation in which a domain of TRAK2 folds back to blocks its kinesin-binding domain (van Spronsen et al., 2013). We, however, observed that both TRAK1 and TRAK2 could recruit kinesin to PEX3Miro1 peroxisomes but that TRAK1 was quantitatively more effective than TRAK2. This observation aligns with the differences previously found between TRAK1 and TRAK2 but also suggests that, in a cellular context, TRAK2 is not exclusively in its closed state and can also support +-end directed mitochondrial transport by KIF5C.

We were surprised to find that PEX3Miro2 failed to recruit either TRAK, KIF5C, or P135 to peroxisomes and that this failure could not be reversed by mutations of the GTPase domains. We do not think this a consequence of the tags at the N-terminal because a similarly tagged form of Miro2 was shown to bind Myo19 (Bocanegra et al., 2020). It is possible that Miro2 requires additional factors present on mitochondria, such as Metaxins, that were not on peroxisomes, but it is also possible that Miro2 does not mediate mitochondrial transport by microtubule-based motors. Miro2 has received less attention than Miro1 and there is less evidence for a role of Miro2 in driving long-range mitochondrial motility. Loss of Miro2 had little effect on mitochondrial motility in hippocampal neurons (Lopez-Domenech et al., 2016). The persistence of some processive mitochondrial movements in cells lacking Miro1 and the severe phenotypes of Miro1 and 2 double knockouts (Lopez-Domenech et al., 2016; Nguyen et al., 2014) has suggested some redundancy in their function. Although overexpressed Miro1 and Miro2 both localize to all mitochondria (Fransson et al., 2003), it is not known whether this is true of the endogenous proteins or if they localize to different populations of mitochondria or different sites on mitochondria. Myo19 does interact with Miro2 to support actin-based mitochondrial transport (Oeding et al., 2018) and Miro2 is implicated in tethering mitochondria to ER (Modi et al., 2019; White et al., 2020). Thus, the sharp difference in the behavior of PEX3Miro1 and PEX3Miro2 reported here indicates that further attention to their functional differences is needed.

We note that our data disagree with that of (Fransson et al., 2006) who studied overexpression of Miro and TRAK and found that TRAK coprecipitated with both Miro1 and Miro2 and did so regardless of the state of the N-GTPase. We propose that this was due to the coassembly of the overexpressed forms of Miro with the endogenous wildtype Miro1 which in turn bound to TRAK.

GTPases are regulated by GTPase-activating proteins (GAPs) and Guanine-nucleotide exchange factors (GEFs) and a Miro GAP and GEF would be expected in cells if the GTPase domains are behaving in vivo as regulatory switches. VopE, a protein expressed by *Vibrio Cholera* Type III in infected cells behaves as a GAP for Miro. By converting the N-GTPase of both Miro1 and Miro2 to the GDP-state it alters the distribution of the host cell’s mitochondria in a manner consistent with our findings (Suzuki et al., 2014). Our data would indicate that VopE will do so by causing the dissociation of TRAK from Miro and thereby releasing kinesin and dynein from the mitochondrial surface. The activity of a GAP from a pathogen begs the question as to what endogenous cellular proteins function as Miro GAPs and when they are called upon to disassemble the mitochondrial motor-adaptor complex. No such Miro GAPs are known at present. Vimar, however, the *Drosophila* homolog of RAP1GDS1, may be a potential Miro GEF; Vimar mutations alter mitochondrial morphology in *Drosophila* and RAP1GDS1 knockdown rescues damaged mitochondrial phenotypes in mammalian cells (Ding et al., 2016). RAP1GDS1 coprecipitates with Miro1 but it has not been shown to alter the GTP/GDP state of Miro as predicted for a GEF. GBF1, a GEF for Arf1 GTPase (Gillingham and Munro, 2007), also influences mitochondrial morphology and position (Walch et al., 2018) and is a candidate for a cellular Miro GEF. Changes in mitochondrial morphology and distribution, however, can reflect many factors beyond the activity of the microtubule-based motors, including the fusion and fission apparatuses and mitochondrial contacts with other organelles. The PEX3Miro system may be valuable for identifying cellular GEFs and GAPs that control Miro1 and thereby control mitochondrial movement.

From our imaging of peroxisomes in neurons as well as COS-7 cells, it is clear that the state of the N-GTPase can have a profound effect on organelle distribution. These changes in distribution are consonant with a model in which the N-GTPase is a switch that controls TRAK, kinesin, and P135 assembly into a Miro1-based motor-adaptor complex. We do not know when in a neuron the N-GTPase is in the GDP state. In neurons expressing labeled kinesin, all the axonal mitochondria, including those that are stationary, have kinesin on them (Wang and Schwarz, 2009) and some signals that trigger mitochondrial arrest, including elevated Ca^2+^ or glucose, do not cause kinesin to fall off mitochondria (Basu et al., 2021; Wang and Schwarz, 2009); these signals cannot be acting through the N-GTPase. Motors are shed from mitochondria, however, during metaphase in mitotic cells (Chung et al., 2016) and it is possible that this is accompanied or mediated by a change in the GTP state of Miro. Understanding the mechanism by which the GTPase switch influences the complex, as revealed by the present studies, should facilitate elucidating when and how cells use this switch to control mitochondrial behavior.

## Materials and methods

### Primers

Sequences of all primers used in this study are presented in Table S2

### Plasmid Constructs

#### Previously Published constructs

The following previously published DNA constructs were used in this study: pmTurquoise2-Peroxi was a gift from Dorus Gadella (University of Amsterdam (Addgene plasmid # 36203; http://n2t.net/addgene:36203; RRID:Addgene_36203) (Goedhart et al., 2012), pmTurquoise2-Mito was a gift from Dorus Gadella (University of Amsterdam (Addgene plasmid # 36208 ; http://n2t.net/addgene:36208 ; RRID:Addgene_36208) (Goedhart et al., 2012), mCitrine-Peroxisomes-2 was a gift from Michael Davidson (National MagLab (Addgene plasmid # 54672; http://n2t.net/addgene:54672 ; RRID:Addgene_54672), mCitrine-C1 was a gift from Robert Campbell & Michael Davidson & Oliver Griesback & Roger Tsien (Howard Hughes Medical Institute, University of California, San Diego and National MagLab (Addgene plasmid # 54587; http://n2t.net/addgene:54587 ; RRID:Addgene_5487) (Griesbeck et al., 2001)). mycMiro1-V13, V427, N18, N432 were kindly provided by Dr. P. Aspenstrom (Karolinska Institute). The construction of Myc-TRAK1 and PEX3-6xhis-mRFP-Miro1 in our lab was previously described (Basu et al., 2021; Pekkurnaz et al., 2014).

#### Purchased Constructs

mCitrine-P135 was subcloned by Vector Builder from an mCitrine-P150Glued

#### Constructs cloned in this study

1. To make the PEX3-6xhis-mRFP construct (PEX3-C), the PEX3-6xhis-mRFP-Miro1 construct (PEX3Miro1) described in Basu et al., 2021 was linearized by PCR using Primers *KP14* and *KP15* to amplify the PEX3-6xhis-mRFP segment of the plasmid and the plasmid backbone but omit Miro1. The PCR product was ligated using the KLD reagent from the Q5 site-directed mutagenesis kit from NEB. This construct was used as a control throughout all co-localization experiments.
2. To make the myc-TRAK2 construct, the Myc-TRAK1 construct described in (Pekkurnaz et al., 2014) was linearized by PCR with the primers *KP51* and *KP52* thereby excluding the TRAK1 sequence from a vector still containing the myc-sequence. A TRAK2 cassette was amplified from a TRAK2 cDNA (BC048093, Transomic) using *KP49* and *KP50* primers that had appropriate overhangs to the myc- vector for recircularization by Gibson assembly. This construct was used to overexpress myc-TRAK2 in kinesin colocalization experiments.
3. mCitrine-KIF5C-YFP was derived from the mCitrine-KHC-eCFP construct generously gifted by Kristen Verhey (University of Michigan (Cai et al., 2007)). The eCFP tag was substituted with a YFP tag.
4. To make the mCitrine-TRAK1-YFP and mCitrine-TRAK2-YFP constructs, the mCitrine-KIF5C-YFP was linearized by PCR with the primers *KP61* and *KP62* for TRAK1 and *KP65* and *KP66* for TRAK2. These primers were designed to omit the KIF5C from the backbone while retaining the mCitrine-YFP backbone sequence. The TRAK1 and TRAK2 cassettes were amplified from the myc-TRAK1 construct (Pekkurnaz et al., 2014) and myc-TRAK2 construct described in this paper using *KP63* and *KP64* for TRAK1 and primers *KP67* and *KP68*. The primers had appropriate overhangs to the mCitrine-YFP vector for recircularization by Gibson assembly after incorporating the TRAK1 and the TRAK2 cassettes downstream of mCitrine. These constructs were used in all mCitrine-TRAK1 or 2 colocalization experiments.
5. The PEX3-6xhis-mRFP-Miro2 construct was made by amplifying the first 592 amino acid residues from of human Miro2 as cloned in (Brickley and Stephenson, 2011; Fransson et al., 2003; Fransson et al., 2006; Glater et al., 2006; Stowers et al., 2002) with the primers *<u>pex-miro2_ForPrimer</u>* and *<u>pex-miro2_RevPrimer</u>*. The backbone containing PEX3-6xhis-mRFP was amplified from the PEX3-6xhis-mRFP-Miro1 using the primers *<u>pexBackbone_ForPrimer</u>*and *<u>pexBackbone_RevPrimer.</u>* The two PCR fragments were then annealed by Gibson assembly. All the mutations were made by site-directed mutagenesis carried out on the PEX3Miro1 and PEX3Miro2 constructs.
6. To generate the mTurquoise-KIF5C construct, the previously described mCitrine-KIF5C-YFP plasmid, was used to amplify KIF5C using the *KP69* and *KP70* primers that had appropriate overhangs for the mTurquoise vector. The mTurquoise plasmid vector was amplified from pmTurquoise2-Peroxi (Addgene plasmid # 36203 (Goedhart et al., 2012)) using the *KP71* and *KP72* primers. The linearized mTurquoise-vector was re-circularized after incorporating the KIF5C cassette at the C-terminus of mTurquoise by Gibson assembly. These constructs were used to overexpress mTurquoise-KIF5C in colocalization experiments.
7. All PEX3Miro1 GTPase mutants were made using the Q5® Site-Directed Mutagenesis Kit from New England Biolabs. The T18N mutation was made using *KP24* and *KP25*. The P13V mutation was made using *KP26* and *KP27*. The K427N mutation was made using *KP146* and *KP147*. The S432N mutation was made using *KP31* and *KP32*. For mutations where both the N and C-GTPases were mutated we used the N-GTPase mutant as a template for introducing the C-GTPase mutation with the primers mentioned above.
8. All PEX3Miro2 GTPase mutants were made using the Q5® Site-Directed Mutagenesis Kit from New England Biolabs. The T18N mutation was made using *KP78* and *KP79*. The A13V mutation was made using *KP76* and *KP77*. The A425V mutation was made using *KP82* and *KP83*. The S430N mutation was made using *KP102* and *KP81*.
9. To make mCitrine-P150Glued we utilized the mCitrine-C1 cloning vector (Griesbeck et al., 2001) and digested the vector with HindIII and Kpn1 enzymes. The P150-Glued cassette was amplified from the pEGFPC2-p150Glued construct provided by Dr. E.L. Holzbaur (University of Pennsylvania) using primers KP112 and KP113 with overhangs with HindIII and Kpn1 enzyme cut sites. The amplified cassette was digested by HindIII and Kpn1 enzymes and the mCitrine-C1 vector and P150 insert were ligated using T4 DNA ligase (NEB BioLabs).

### Antibodies used for western blotting

For Western blots, the following primary antibodies were used at the stated dilutions: anti-human TRAK1 at 1:2000 (HPA005853, Sigma-Aldrich), anti-myc at 1:5000 (05-724, EMD Millipore), anti-RFP at 1:1000 (SAB2702214, Sigma-Aldrich) and anti-6X-HIS at 1:1000 (MA1-21315, Thermo Fisher Scientific). For fluorescent detection (used for all quantitative blots) 680RD Donkey anti-Rabbit, 800CW Donkey anti-Mouse were used at 1:5000 (LiCor Biosciences) and all blots were scanned by the Odyssey® CLx Imaging System.

### Cell Cultures and transfections

HEK293T and COS-7 cells were cultured in DMEM supplemented with L-glutamine, penicillin/streptomycin (Life Technologies), and 10% FBS (Atlanta Premium). Plasmid DNA transfections in HEK293T cells were performed with the calcium phosphate (Kingston et al., 2003). Plasmid DNA transfections in COS-7 cells were performed with PolyJet DNA (Signa Gen Laboratories) using the manufacturers guidelines. These cell lines were generally transfected 18-24 hours after plating and fixed 2 days later.

**Hippocampal neurons** were dissected and dissociated from E18 rat (Charles River) embryos as previously described (Nie and Sahin, 2012) and plated at 5–7 × 10^4^/cm^2^ density on glass bottom dishes (D35-20-1.5-N, Cellvis) coated with 20 μg/ml poly-L-Lysine (Sigma) and 4 μg/ml laminin (Life Technologies) and maintained in neurobasal (NB) medium supplemented with B27 (Life Technologies), L-glutamine, and penicillin/streptomycin, unless specified otherwise. Hippocampal neurons were transfected on DIV5 using Lipofectamine2000 (11668-019, Life Technologies) and imaged 2–3 days later.

### Immunoprecipitations

For all immunoprecipitations, HEK293T cells were plated at 5.5×10^5^ cells/well density in a 6-well plate and transfected the next day. Two days after transfection, cells were washed once with ice-cold 1X phosphate-buffered saline (1XPBS: (NaCl: 137 mM, KCl: 2.7 mM, Na_2_HPO_4_: 10 mM, KH_2_PO_4_: 1.8 mM)) and lysed in 600μl PierceTM IP Lysis Buffer (Thermo Fisher, Catalog number: 87787) and protease inhibitor cocktail set III (539134-1SET, EMD Millipore) at 1:500 dilution. Lysates were centrifuged 10 min at 13,000 x g at 4°C and the clarified supernatants were collected. For immunoprecipitations of PEX3-6xhis-mRFP-Miro, anti-6X-HIS antibody was incubated for 1 hr at 4°C with 500 μl of the clarified supernatants of whole-cell lysates and then for 1 more hour with Dynabeads Protein G beads (Thermo Fisher, catalog number: 10003D). The beads had been pre-blocked in .2% TBST and washed three times with lysis buffer. After magnetic separation, beads were washed 5 times in lysis buffer and resuspended in 2xLaemmli buffer. 50% of this fraction was then separated by SDS-PAGE and transferred to nitrocellulose membranes. The membranes were pre-blocked overnight with 3% bovine serum albumin (w/v) in 1X Tris-buffered saline (TBS) with 0.1% (w/v) Tween20 (TBS: 20 mM Tris, NaCl: 150 mM), followed by overnight incubation at 4C with primary antibodies in the blocking buffer. The blot was then washed 3 times with TBST and blotted with secondary antibodies for 1 hour at room temperature, before ECL imaging or being scanned by the Odyssey® CLx Imaging System.

### Imaging Acquisition and Quantification

For imaging co-localization of TRAK1, TRAK2, P135, and KIF5C on PEX3Miro1 peroxisomes, COS-7 cells were grown on glass bottom dishes (FD35-100, World Precision Instruments, Sarasota County, FL, United States of America). Cells were fixed and imaged at room temperature. For imaging the distribution of PEX3Miro1 peroxisomes in rat hippocampal neurons, primary neurons were isolated as described and grown on glass bottom dishes (FD35-100, World Precision Instruments, Sarasota County, FL, United States of America). Neurons were fixed 48 hours after transfection and imaged at room temperature. COS-7 cells and neurons were imaged on a Leica SP8 laser scanning confocal microscope. Images were acquired using a 60X objective. A White Light Laser and Argon laser were used at 70% laser intensity and line scanning with 3 scans per line was used to acquire all images. The percentage of laser intensity used for the 458 spectra to capture mTurquoise markers was 5% in all experiments. The percentage of laser intensity used for the 515 spectra to capture mCitrine-tagged constructs was 10%. The percentage of laser intensity used for the 554 spectra to capture all mRFP tagged constructs was 10%. Gain was set at 100 for imaging mCitrine-TRAK1, mCitrine-TRAK2, mCitrine-KIF5C, PEX3-C, and PEX3Miro for all COS-7 and hippocampal neuron images. Only cells with visible PEX3-C or PEX3Miro signals on peroxisomes at 100 gain with these laser settings were imaged to minimize differences in expression. Images were acquired using z-stacks with 0.3μm slices for COS-7cells and 0.2μm for neurons, ranging from the top to the bottom of the cell or neuronal soma.

Images for figure S7 were obtained through the Leica DMi8 thunder microscope system with an sCMOS camera having a pixel size of 6.5uM (DFC 9000) and a 40X 0.8NA objective. Following image acquisition, images were processed through the small volume computational clearing pipeline, proprietary to the thunder imaging system. Each neuron was imaged as a series of areas later stitched together to visualize the entire neuron. In addition to imaging in single channels, neurons were also illuminated simultaneously for CFP (mTurquoise-SRL) and YFP (mCitrine-KIF5C-YFP) fluorescence to obtain images that represent the whole neuron (cell fill).

### Quantifications of organelle distribution in COS7 cells

Quantification of peroxisome distribution was done as described in (Basu et al., 2021) using the DoveSonoPro software (https://github.com/ThomasSchwarzLab). DoveSonoPro software allows for the cell outline and cell center to be selected by the user. Image file names were blinded for quantification.

### Colocalization Quantification

The co-localization of TRAK1/2, KIF5C, and P135 with PEX3-C or PEX3Miro peroxisomes was quantified using a custom FIJI macro. Images of cells were acquired using z-stacks with .3um thickness and z-slice images were acquired from top to the bottom of cells to account for all peroxisomes. This custom macro creates a mask of the peroxisome marker channel (mTurquoise-SRL or mCitrine-PTS1) in each z-slice and uses the peroxisome mask to analyze only the pixels in the image where the peroxisome marker co-localizes with the RFP-tagged PEX3-C or PEX3Miro construct. The PEX3-C and PEX3Miro positive peroxisomes are then analyzed for co-localization KIF5C, TRAK1, TRAK2, P135. The final quantification is the average of the raw enrichment of KIF5C, TRAK1, TRAK2, or P135 across z-slices on PEX3-C or PEX3Miro1 positive peroxisomes. Because the expression of the PEX3 control or PEX3Miro1 construct varies slightly from cell to cell, we measured the intensity of the PEX3-C and PEX3Miro construct that is localized to peroxisomes. This macro can control for any PEX3-C or PEX3Miro1 expression-dependent differences in co-localization by normalizing the raw amount of KIF5C, TRAK1, TRAK2, or P135 to raw intensity of PEX3-C or PEX3Miro expression. We controlled all experiments by ruling out any expression dependent differences. We then control for potential error in co-localization quantification by taking into consideration the partial cytosolic localization of any overexpressed mCitrine-KIF5C, TRAK1, TRAK2, or P135. To do this, we subtract the whole cell intensity of the mCitrine signal that exists outside of the peroxisome mask from the specific signal intensity that overlaps with the peroxisome mask. This background fluorescence subtraction allows for the quantification of the specific amount of signal on PEX3-C or PEX3Miro peroxisomes without cofounding results with the presence of the cytosolic localization of the overexpressed constructs.

### Statistical Analysis

Statistical analysis was performed with GraphPad Prism v7.0. A two-tailed unpaired t-test with Welch correction or a one-way ANOVA with Dunett T3 correction was used to determine significances or differences between populations in all co-localization experiment quantifications. P-values appear in all the Figures. All colocalization quantifications are depicted as Tukey boxplots.

## Summary of Supplementary Materials

### Supplementary Figures

**Figure S1 (related to Figure 1)** shows and quantifies the expression of PEX3-C, PEX3Miro1, and PEX3Miro2 on peroxisomes and mitochondria. **Figure S2 and Figure S3 (related to Figure 5)** show how PEX3Miro1 C-GTPase mutants and double mutants affect the recruitment of mCitrine-TRAK1 and mcitrine-TRAK2 to peroxisomes. **Figure S4 (related to Figure 6)** shows the recruitment of mCitrine-KIF5C to peroxisomes when overexpressed with PEX3Miro1 N-GTPase mutants and myc-TRAK2. **Figure S5 (related to Figure 6)** shows the quantification of the amount of mTurquoise-KIF5C recruited to peroxisomes in the presence of overexpressed mCitrine-TRAK1 or mCitrine-TRAK2. **Figure S6 (related to Figure 7)** shows and measures the expression of mCitrine-TRAK1 and mCitrine-TRAK2 on peroxisomes that have PEX3Miro2 or PEX3Miro2 GTPase mutants. **Figure S7** shows a whole neuron image of peroxisomes marked by mTurquoise-SRL in hippocampal rat neurons.

## Supplementary Tables

**Table S1 (related to methods)** is the list of primer sequences used in this study.

## Acknowledgements

We thank L. Mkhitaryan and Zerong Cai for technical support and the past and present T. L. Schwarz laboratory members for their considerations and critical reading of the manuscript. We thank the Harvard NeuroDiscovery Center’s Enhanced Neuroimaging Core (NINDS P30 Core Center grant no. NS072030) and the Cellular Imaging Core IDDRC at Boston Children’s Hospital (NIH U54 HD090255) for support with imaging. This research was supported by the National Institutes of Health grants R01 GM069808 (NIH/NIGMS) to T.L.S. and F31 GM126681-02 (NIH/NIGMS) to K.D.

## Author Contributions

H.B., T.L.S., and K.D. conceived the project. H.B. constructed the PEX3-C, PEX3Miro1, and PEX3Miro2 constructs and created the co-localization quantification code as well as the previously published DoveSonoPro code. E.S. helped optimizing experiments with mCitrine-TRAK1, mCitrine-TRAK2, and mTurquoise-KIF5C and assisted in analyzing peroxisome distribution. K.D. performed all other experiments. T.L.S. and K.D. wrote the manuscript with feedback from all authors.

## Supplementary Information

**Figure S1:**
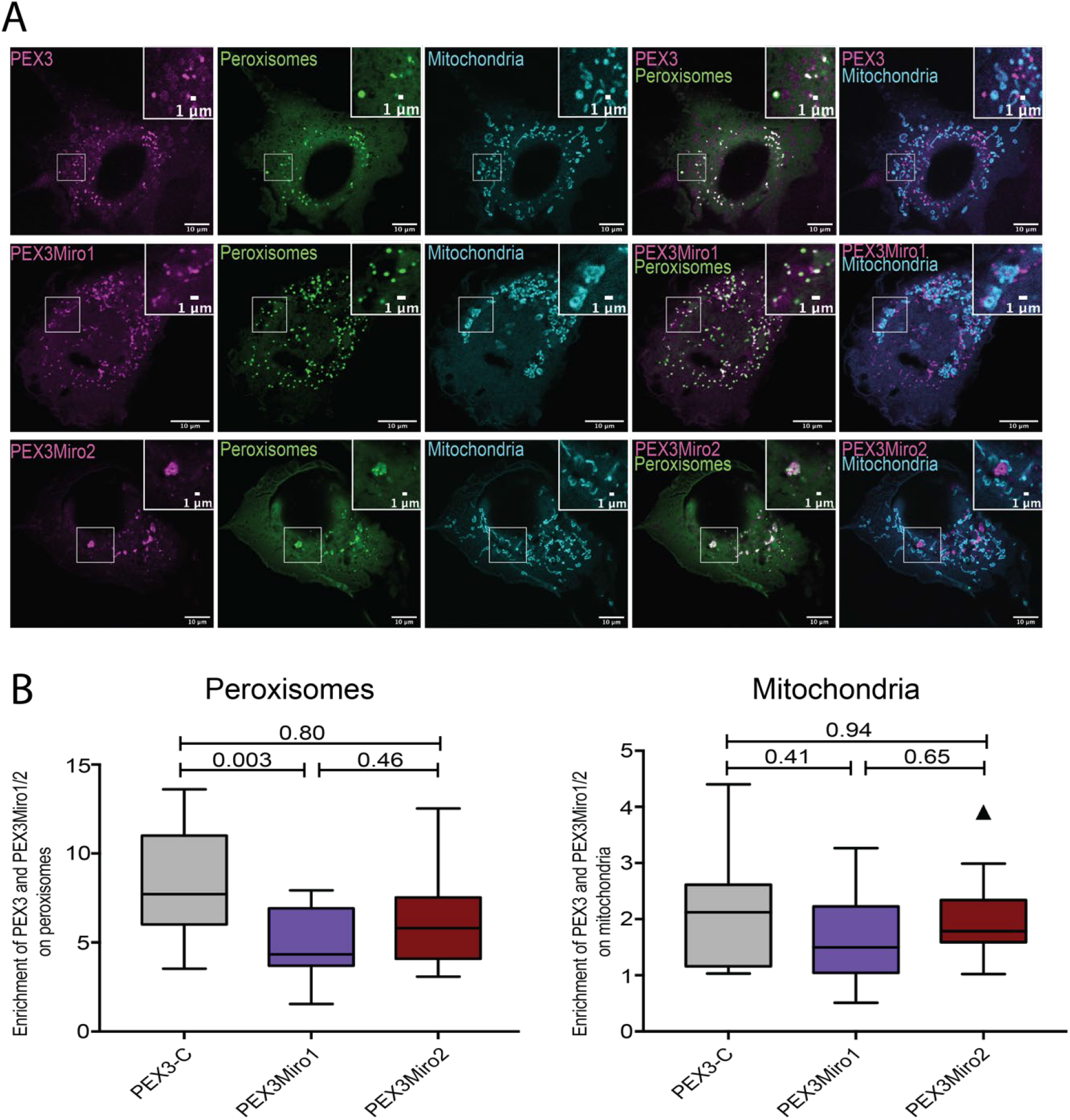
PEX3, PEX3Miro1 and PEX3Miro2 expression on peroxisomes and mitochondria. A) COS-7 cells were co-transfected with PEX3-C, PEX3Miro1, or PEX3Miro2 together with the peroxisome marker mCitrine-PTS1 (green) and the mitochondrial marker Mito-mTurquoise (blue) N=15 cells over 3 biological replicates. The corner inserts show enlargement of the boxed regions. B) From cells as in (A), quantification of the RFP tags on PEX3-C and PEX3Miro that colocalized with the peroxisomal and mitochondrial markers. The data are represented as ‘Box and Whiskers’ plots with the median values indicated. The indicated P values were obtained from One way ANOVA with Dunnett’s T3 correction for multiple comparisons. N=15 cells over 3 biological replicates.

**Figure S2:**
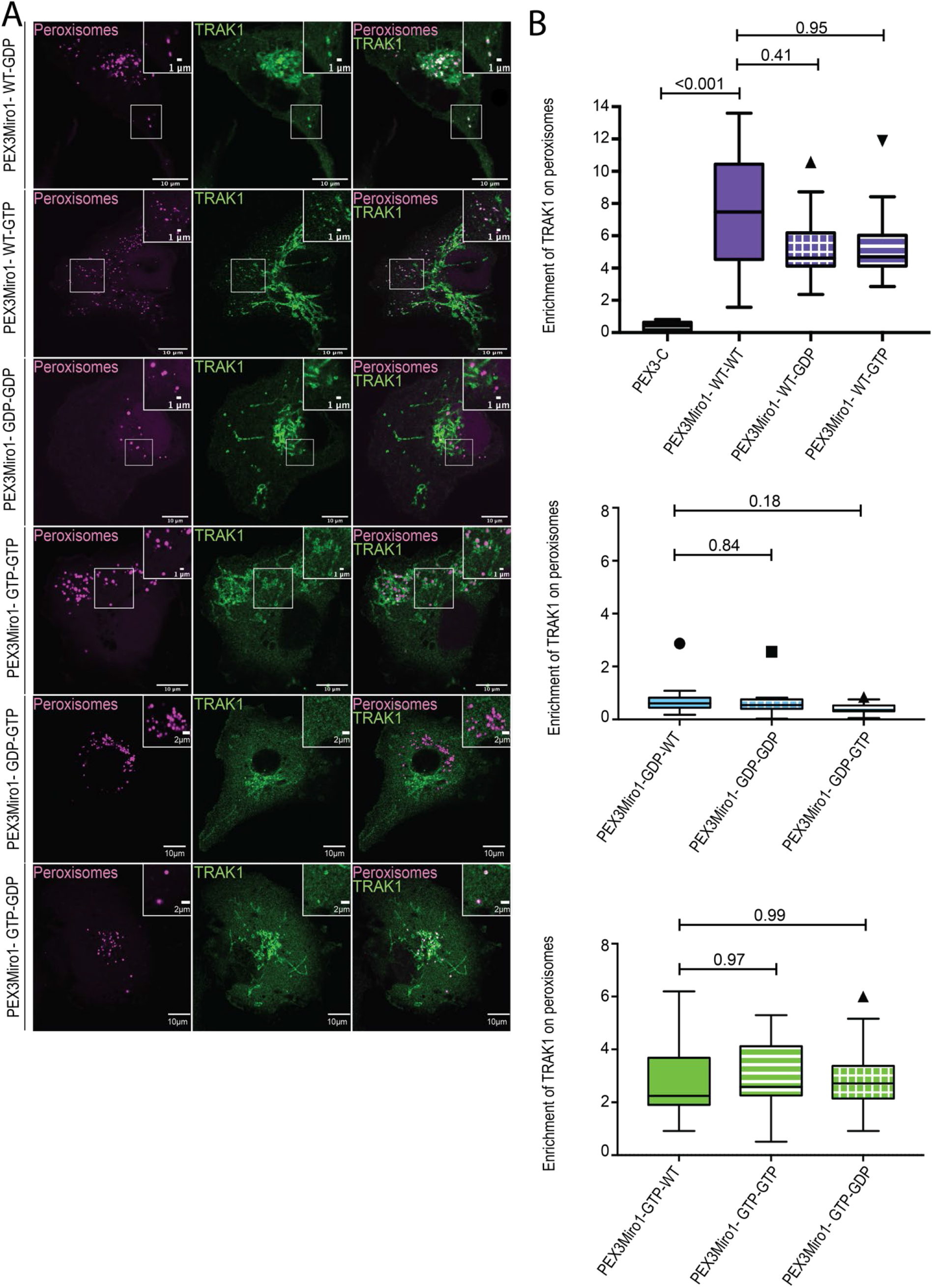
The PEX3Miro1 C-GTPase has little influence on the co-localization of TRAK1 and PEX3Miro1. A) PEX3Miro1, with the indicated mutations of the N- and C-GTPase domains, was coexpressed with mCitrine-TRAK1 (green) and mTurquoise-SRL (magenta) in COS-7 cells. The corner inserts show enlargement of the boxed regions. B) Quantification of the amount of TRAK1 enriched on peroxisomes transfected as in (A). The data are presented as ‘Box and Whiskers’ plots with the median value indicated. Outliers are represented as individual dots and are considered in all statistical calculations. The indicated P-values were determined by one-way ANOVA with Dunnett’s T3 correction for multiple comparisons. N=15 cells over 3 biological replicates. All experiments were conducted and imaged in parallel except the GDP-GTP and GTP-GDP forms of PEXMiro1; they were performed together with controls that were consistent with the values of the other data shown.

**Figure S3:**
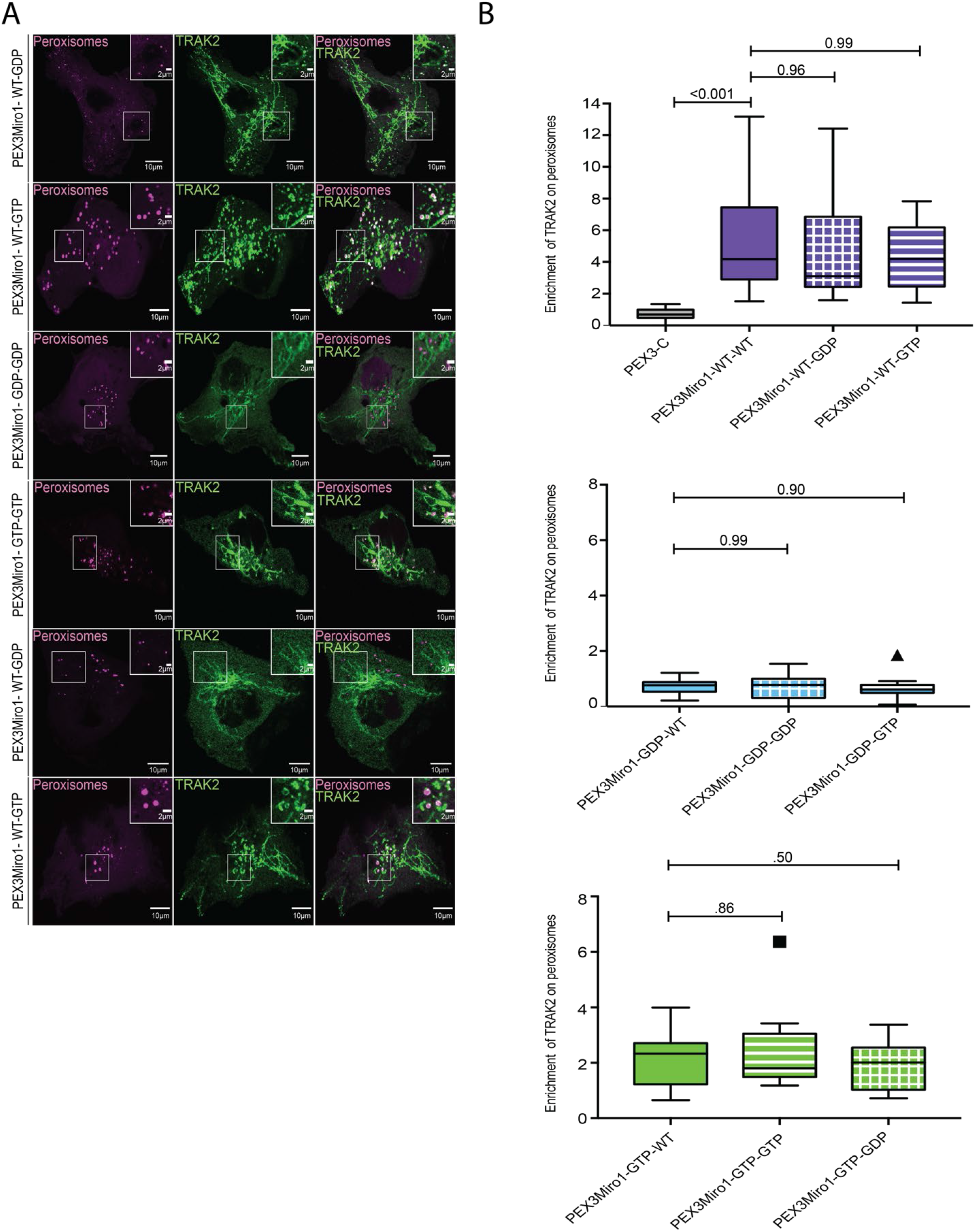
The PEX3Miro1 C-GTPase has little influence on the co-localization of TRAK2 and PEX3Miro1. A) PEX3Miro1, with the indicated mutations of the N- and C-GTPase domains, was coexpressed with mCitrine-TRAK2 (green) and mTurquoise-SRL (magenta) in COS-7 cells. The corner inserts show enlargement of the boxed regions. B) Quantification of the amount of TRAK2 enriched on peroxisomes transfected as in (A). The data are presented as ‘Box and Whiskers’ plots with the median value indicated. Outliers are represented as individual dots and are considered in all statistical calculations. The indicated P-values were determined by one-way ANOVA with Dunnett’s T3 correction for multiple comparisons. N=15 cells over 3 biological replicates. All experiments were conducted and imaged in parallel except the GDP-GTP and GTP-GDP forms of PEXMiro1; they were performed together with controls that were consistent with the values of the other data shown.

**Figure S4:**
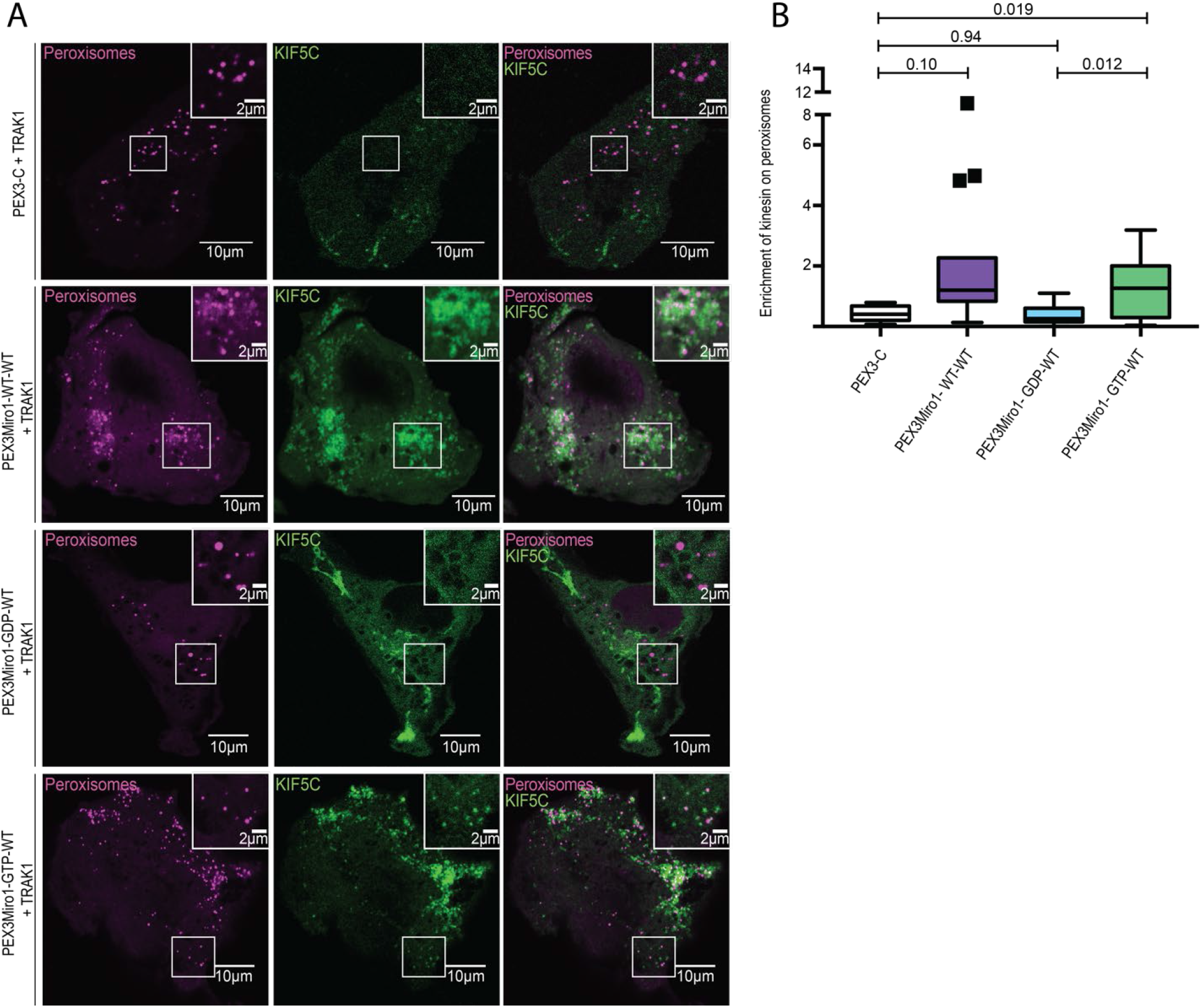
The PEX3Miro1 N-GTPase regulates co-localization of KIF5C in the presence of TRAK2. A) Either wildtype or N-GTPase mutations of PEX3Miro were expressed in COS-7 cells together with myc-TRAK2, mCitrine-KIF5C(green) and mTurquoise-SRL (magenta). Expression of PEX3Miro constructs on peroxisomes was confirmed by visualization of their RFP tags. The corner inserts show enlargement of the boxed regions. B) From cells as in (A), quantification of mCitrine-KIF5C on peroxisomes. The data are presented as ‘Box and Whiskers’ plots with the median value indicated. The indicated P values were determined by one-way ANOVA with Dunnett’s T3 correction for multiple comparisons. Outliers are represented as single points and included in all statistical calculations. N=15 cells over 3 biological replicates

**Figure S5:**
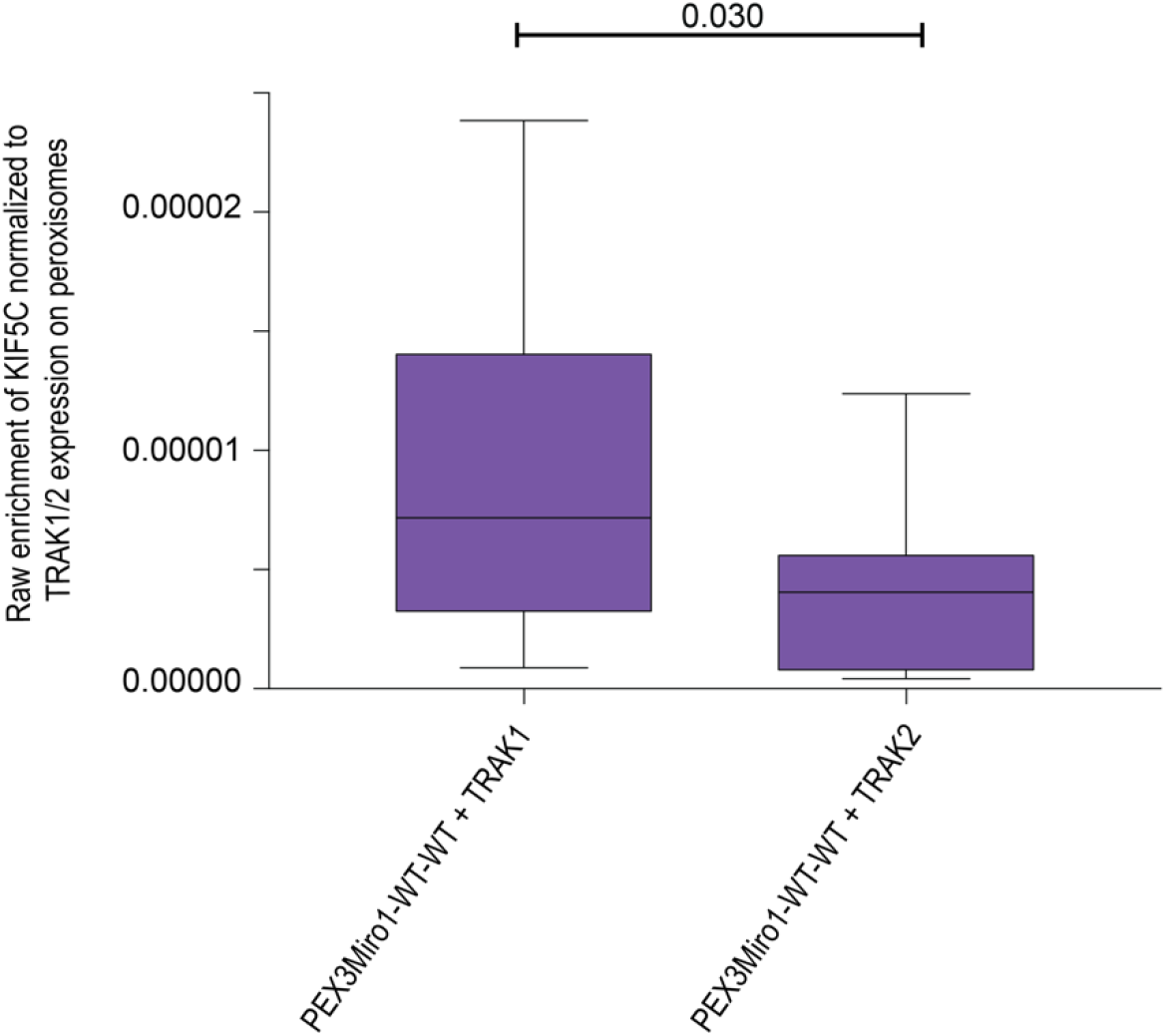
TRAK1 and 2 differ in their recruitment of KIF5C to peroxisomes. COS-7 cells were transfected with mTurquoise-KIF5C, PEX3Miro1, and either mCitrine-TRAK1 or mCitrine-TRAK2. The amount of mTurquoise on peroxisomes was normalized to the intensity of mCitrine on peroxisomes to determine the relative efficacy of the two TRAK isoforms for recruiting KIF5C. The data are presented as ‘Box and Whiskers’ plots with the median value indicated. The indicated P-value was obtained from the two-tailed unpaired t-test with Welch correction. N=15 cells over 3 biological replicates.

**Figure S6.**
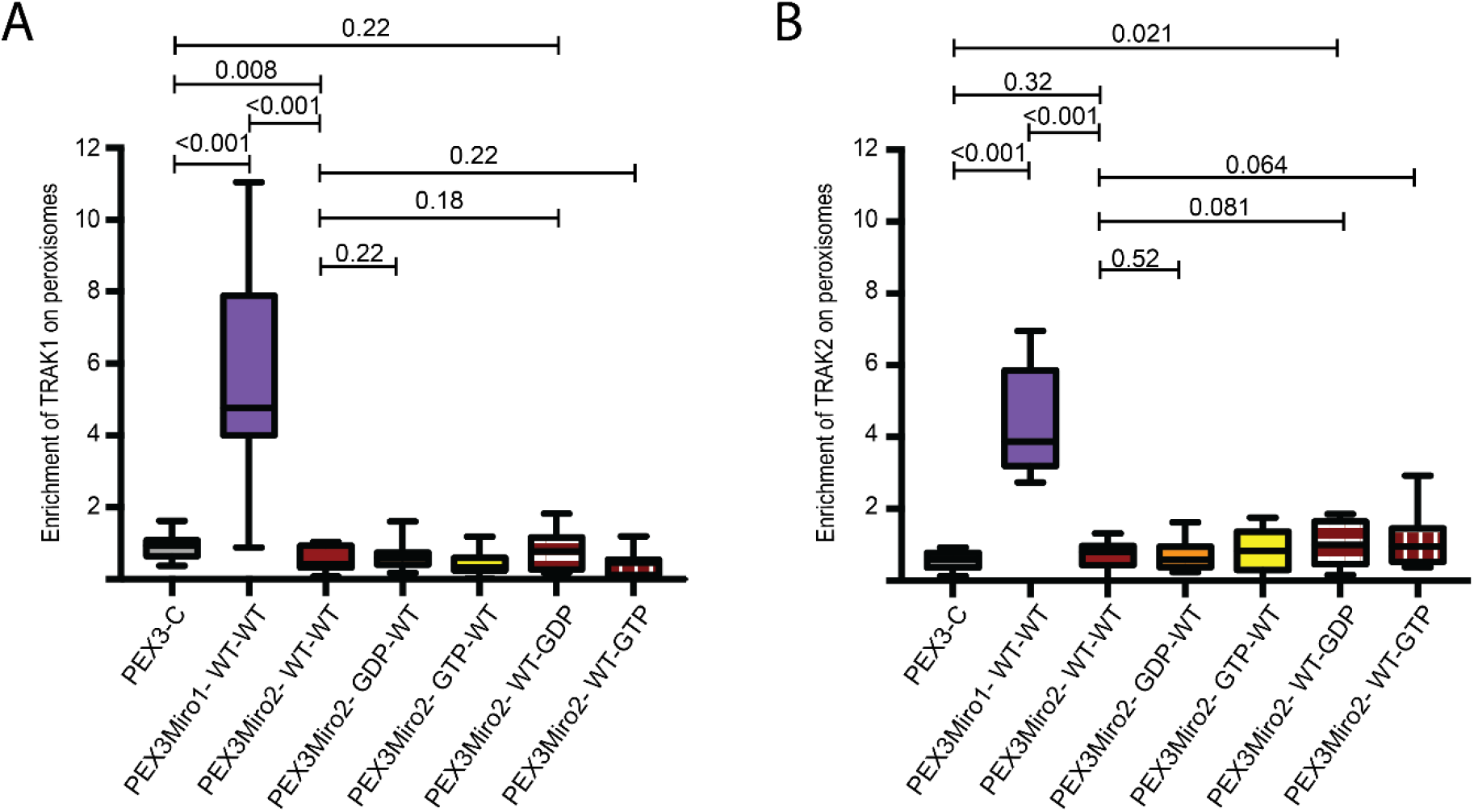
PEX3Miro2 does not recruit TRAK1 or TRAK2 to peroxisomes. Quantification of the amount of mCitrine-TRAK1 (A) or mCitrine-TRAK2 (B) enriched on peroxisomes carrying PEX3-C, PEX3Miro1, PEX3Miro2, or the PEX3Miro2 GTPase mutant constructs as indicated on the x-axis. Peroxisomes were marked with mTurquoise-SRL. The quantification is represented as ‘Box and Whiskers’ plots with the median value indicated. Outliers have been removed from this data set using the ROUT method (Q=1%) and are not included in statistical calculations. Indicated P values were determined by one-way ANOVA with Dunnett’s T3 correction for multiple comparisons. N=15 cells over 3 biological replicates.

**Figure S7:**
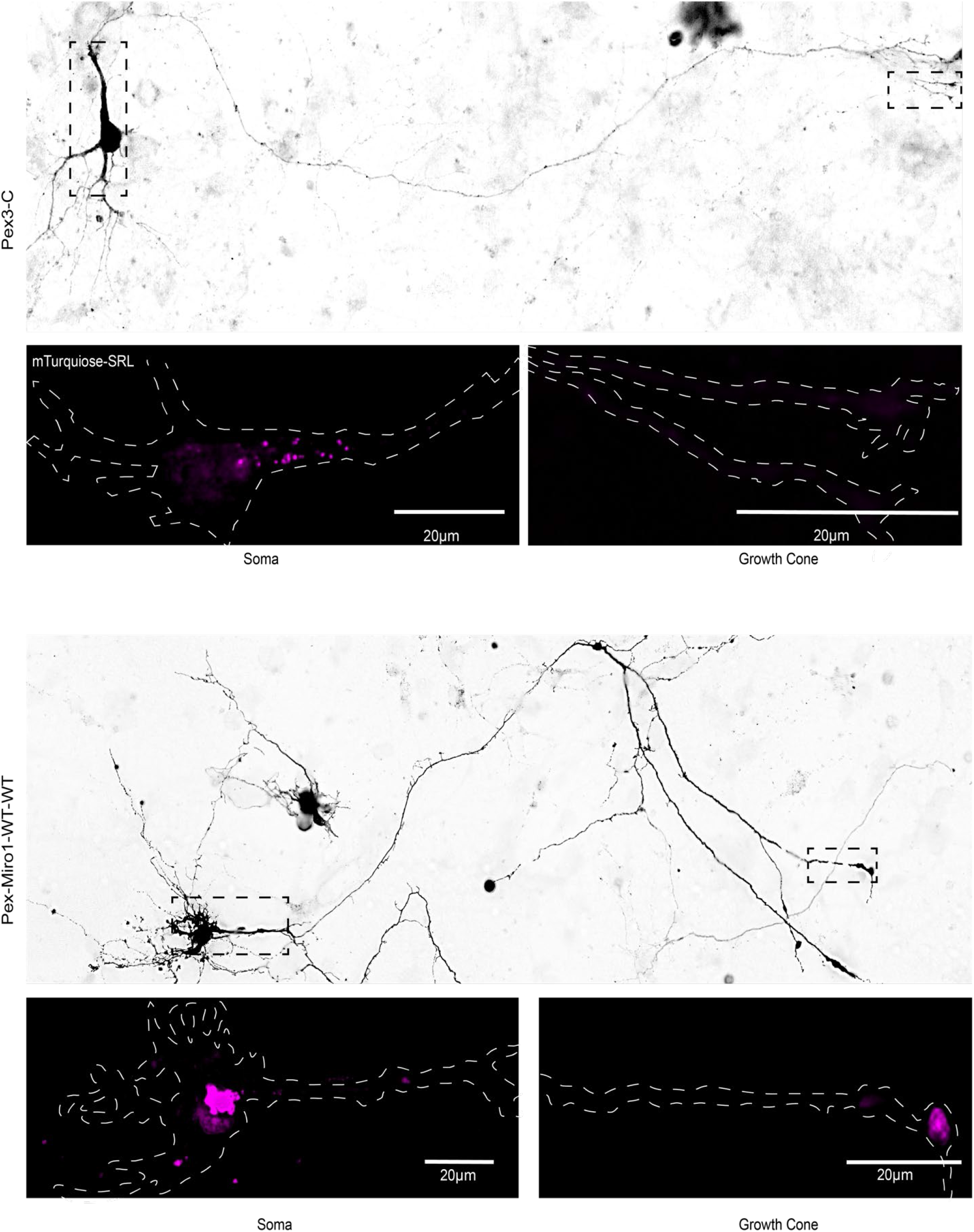
Examples of neurons from which the somatic and growth cone levels of peroxisomes were quantified. Neurons were transfected as in Figure 8, i.e with PEX3-C or a PEX3Miro1 construct, along with mTurquoise-SRL, myc-TRAK1 and mCitrine-KIF5. The mTurquoise-SRL marker was used to visualize the distribution of peroxisomes throughout the neurons, and enabled the peroxisome counts in the soma and growth cone in Figure 7. The mCitrine-KIF5C signal was used to get a low-resolution image of the entire cell. Representative neurons expressing PEX3-C (A) and Pex3Miro1-WT-WT (B) are shown. Upper panels show a full neuron. The soma and growth cone whose peroxisome signals appear below are marked (dotted lines).

**Table S1.**
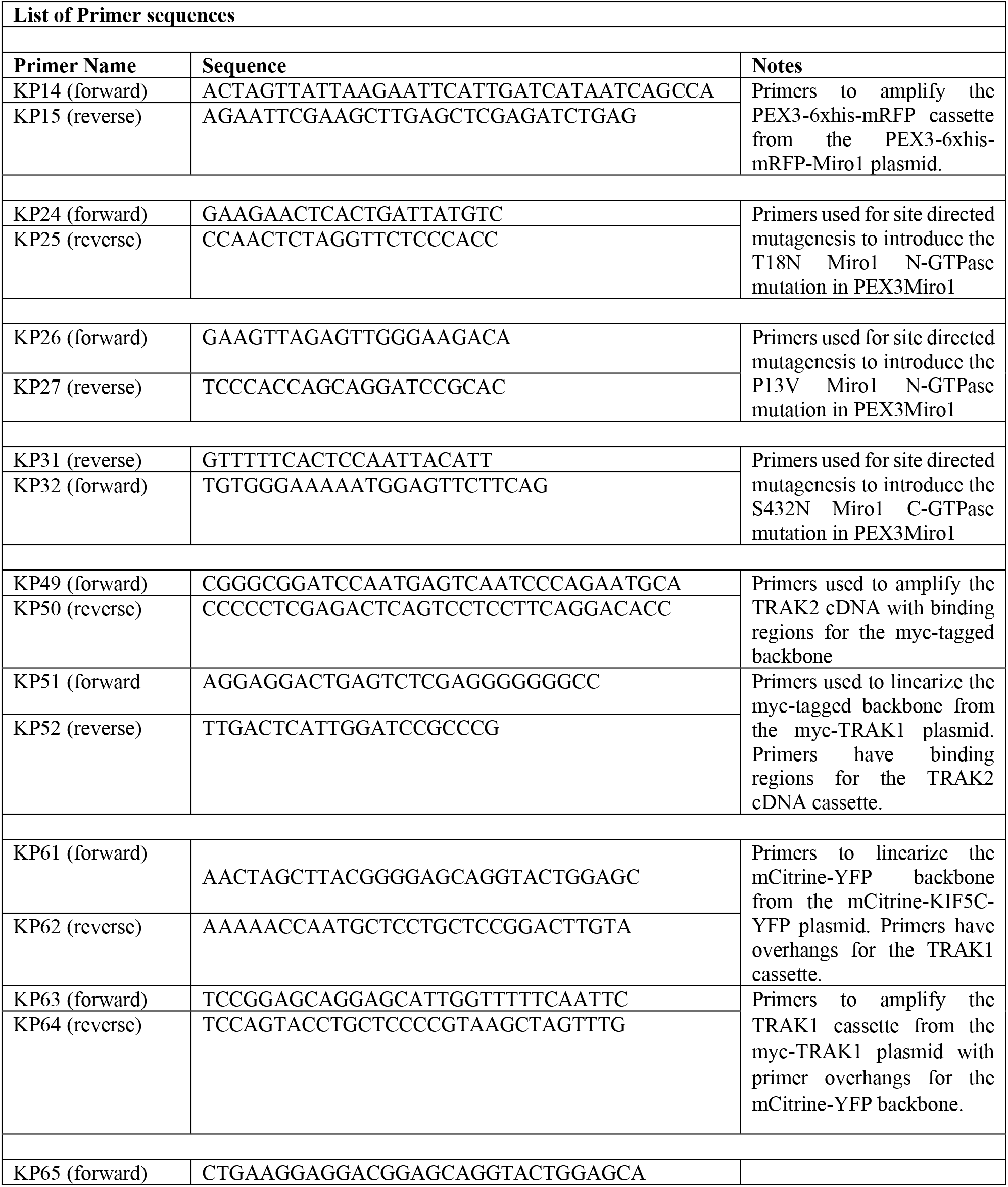
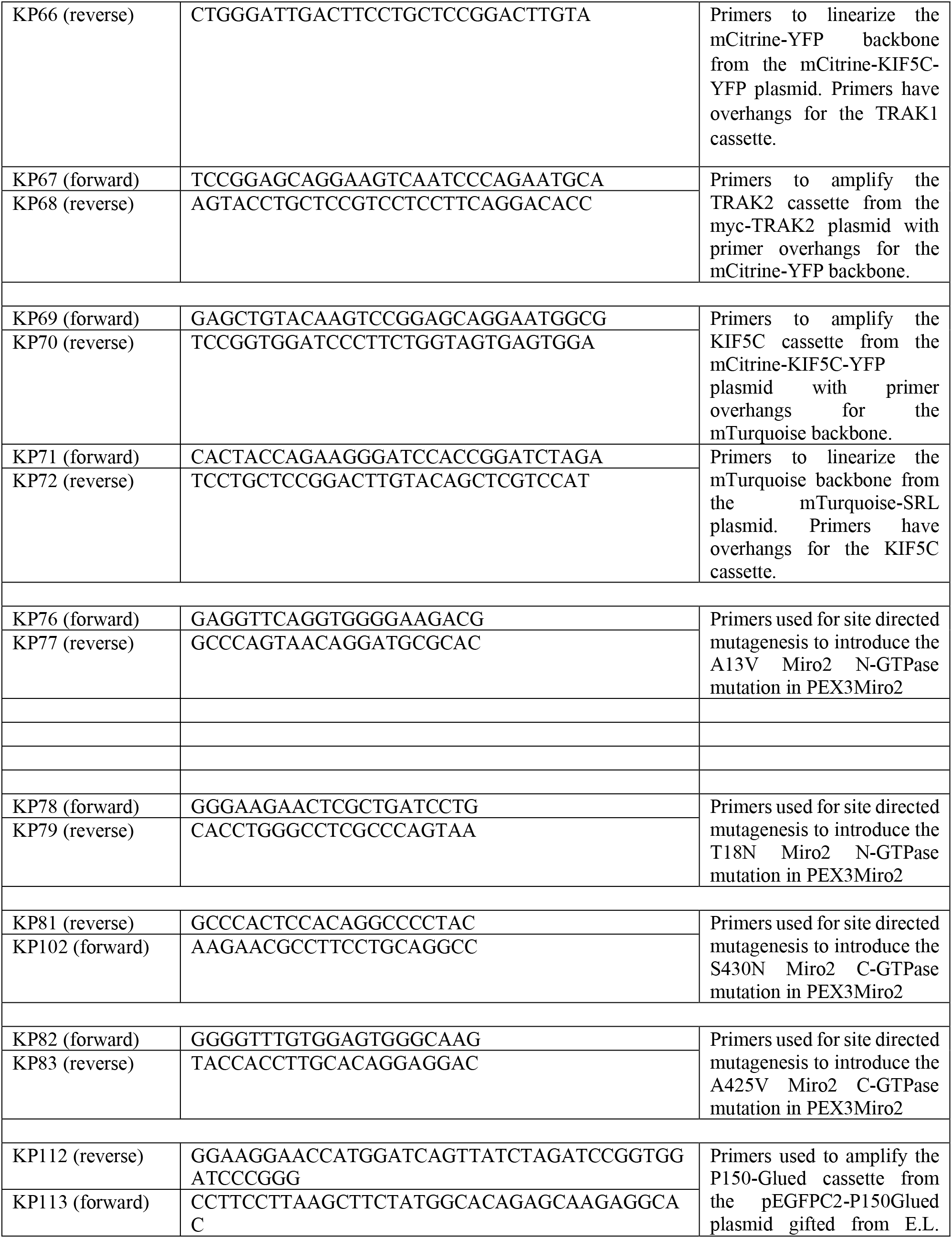
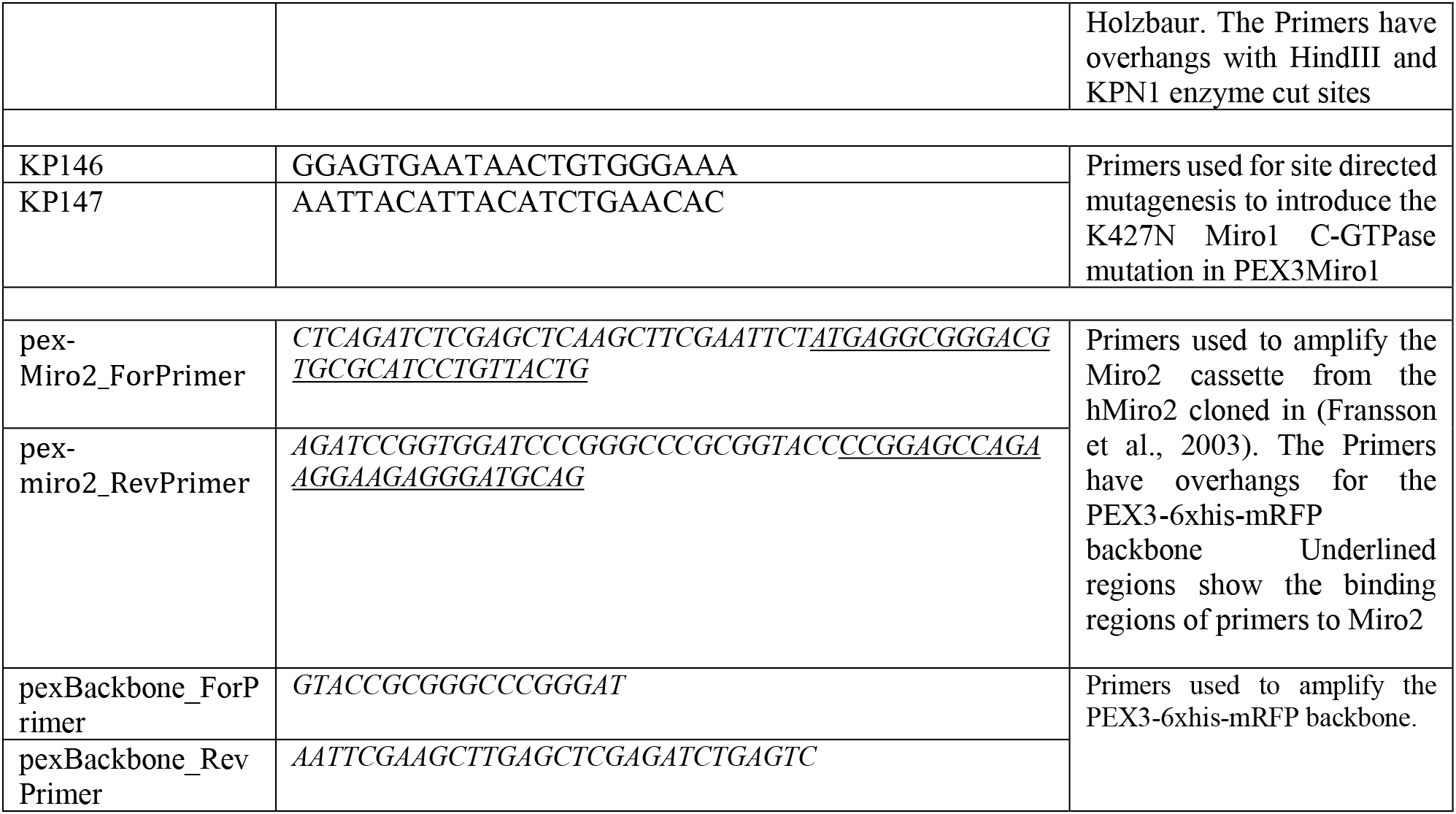
Table of primers used for all plasmids made for this study

